# SIRPA suppresses integrin-dependent virus endocytosis

**DOI:** 10.64898/2026.04.17.719277

**Authors:** Zhonghao Yan, Kruthika Iyer, Minghao Li, Kie Hoon Jung, Ciara Tipping Hu, Natalia Ansin, Nicolás Sarute, Brian B. Gowen, Susan R. Ross

**Author notes:** The 1^st^ two authors contributed equally to this study (Zhonghao Yan, Kruthika Iyer).

## Abstract

New World arenaviruses (NWAs) that cause viral hemorrhagic fever, such as Junín virus, have few therapeutic options. Entry of these viruses into cells is mediated by binding to cell surface receptors, followed by endocytosis and trafficking to a low pH compartment. We showed previously that Signal Regulatory Protein Alpha (SIRPA), a critical cell surface receptor that inhibits macrophage phagocytic activity, decreases internalization by NWAs as well as other pathogenic RNA viruses that traffic to low pH compartments. Here we demonstrate that proteins involved in the SIRPA/integrin signaling axis, including Src homology region 2 (SH2)-containing protein tyrosine phosphatase 2 (SHP2), src family kinases (SFKs), particularly FYN, focal adhesion kinase (FAK), and alpha-integrin play a role in viral endocytosis and that SIRPA inhibits virus entry through blocking this pathway. In addition to defining a role for integrins in viral entry, these studies also provide additional insight into SIRPA’s interference in processes dependent on integrin signaling, including phagocytosis. Moreover, using drugs that block the integrin signaling pathway *in vitro* and *in vivo*, we show that there are additional steps that may be targeted therapeutically for inhibiting infection by RNA viruses that traffic to acidic compartments.

## Introduction

Arenaviruses are enveloped ambisense RNA viruses that infect different species (Emonet et al., 2009; Peralta et al., 1979; Sarute and Ross, 2017). The rodent-borne mammarenaviruses are further categorized into New World (NW) or Old World (OW) arenaviruses based on their geographic endemicity (Charrel and de Lamballerie, 2010). OW arenaviruses originate in the African continent and include Lymphocytic Choriomeningitis virus (LCMV) and Lassa virus (LASV). NW arenaviruses are found in South and North America and are further divided into three clades: A, B, and C (Sarute and Ross, 2017). Most clade B NW arenaviruses, such as Junín virus (JUNV) and Machupo virus (MACV), cause benign infections in their natural rodent hosts but severe disease when they zoonose to humans (Gomez et al., 2011; Lendino et al., 2024); the one exception is Tacaribe virus (TCRV), which was originally isolated from a bat and is likely not highly pathogenic in humans (Bowen et al., 1996; Cogswell-Hawkinson et al., 2012; Schountz, 2024). Clade B NW arenaviruses use human transferrin receptor 1 (TfR1) for cell entry and cause hemorrhagic fever, with up to 30% mortality in humans (Gomez et al., 2011; Lendino et al., 2024; Radoshitzky et al., 2007). Due to the lack of effective treatment, arenaviruses are classified as Category A pathogens by the CDC. Although several inhibitors targeting different stages of arenavirus infection have been reported, there is currently no FDA-approved therapy for arenavirus infection (Iyer et al., 2024). Candid #1, a live attenuated JUNV vaccine strain (JUNV C1), is the only effective arenavirus vaccine in use (Aguilar et al., 2009; Enria et al., 2008). Therefore, new therapeutics are needed.

SIRPA is an N-linked glycosylated transmembrane protein that is highly expressed in myeloid cells but is also found in other cell types (Barclay and Van den Berg, 2014). SIRPA expressed on the surface of phagocytic cells interacts with another transmembrane protein, CD47, found on target cells. This interaction may also occur in *cis* when both proteins are expressed on the same cell (Granda Farias et al., 2025). CD47 expression is highly upregulated on the surface of cancer cells, and CD47/SIRPA intercellular interaction initiates a signaling cascade that dramatically diminishes the ability of cancer cells to be phagocytosed (Barclay and Van den Berg, 2014). Inhibiting SIRPA-CD47 interaction is currently a target of anti-cancer therapies (Bouwstra et al., 2022).

Several players in SIRPA’s upstream and downstream signaling pathway are known. Upon SIRPA/CD47 binding, Src-family kinases (SFK) phosphorylate tyrosines in SIRPA’s cytoplasmic immunoreceptor tyrosine-based inhibitory motif (ITIM) region, which then recruits and activates Src-homology region 2 domain-containing phosphatase-1 or 2 (SHP1/SHP2), leading to suppression of integrin signaling and phagocytosis inhibition (Hayat et al., 2020; Miller et al., 2025; Morrissey et al., 2020). Integrins are heterodimeric, transmembrane receptors consisting of α and β subunits that connect the cell and extracellular matrix to cytoplasmic events by participating in signaling pathways that mediate the reorganization of the intracellular cytoskeleton (Pang et al., 2023). Other components known to function in the integrin signaling pathway include Focal Adhesion Kinase (FAK), Rho proteins and myosin IIA (Alanko and Ivaska, 2016; Clark et al., 1998; Schiller et al., 2013).

We previously showed that SIRPA is a host restriction factor for a number of pathogenic RNA viruses that traffic to acidic endosomes to achieve cell entry (Sarute et al., 2021; Sarute et al., 2019). Both phagocytosis and receptor-mediated endocytosis consist of common steps: receptor binding, membrane invagination, formation of an intracellular vesicle and cytoskeleton rearrangement-dependent vesicle movement to a low pH compartment. Here, we investigated whether molecules involved in SIRPA’s inhibition of phagocytosis are also required for SIRPA’s antiviral activity. We show that similar to SIRPA, SHP2 restricts arenavirus entry, and that tyrosine phosphorylation of SIRPA’s cytoplasmic ITIM domain and SIRPA-SHP2 binding are increased during virus entry; these increases are controlled by the SFK FYN. Since SHP2 activation inhibits integrin-dependent phagocytosis, we tested integrin inhibitors and integrin pathway inhibitors and found that both reduced virus infection, confirming the importance of this pathway in both phagocytosis and infection. The discovery of SIRPA’s antiviral signaling pathway contributes to our understanding of the biology of virus entry and potentially new identifies host targets for antiviral therapeutics for arenaviruses.

## Results

### SHP2 is a virus restriction factor

When SIRPA binds to CD47 on target cells, to surfactants on lung epithelia or after stimulation by growth factors such as EGF, its activation results in phosphorylation of the four tyrosine residues found in its two cytoplasmic ITIMs, followed by the recruitment of SHP phosphatases and inhibition of phagocytosis (Barclay and Brown, 2006). While both SHP1 and SHP2 function in hematopoietic cells, in non-hematopoietic cells, SHP2 carries out this role (Murata et al., 2014). We first tested if SHP2 suppressed virus entry in non-hematopoietic cells. We used RNAi to deplete SHP2 levels in A549 lung cancer epithelial and U2OS osteosarcoma cells, and tested infection by pseudoviruses bearing the influenza type A hemagglutinin, MACV glycoprotein or vesicular stomatitis virus (VSV) glycoprotein (G) protein. In all cases, SHP2 knockdown increased infection to levels similar to that seen with SIRPA knockdown (Fig. 1A).

**Figure 1.**
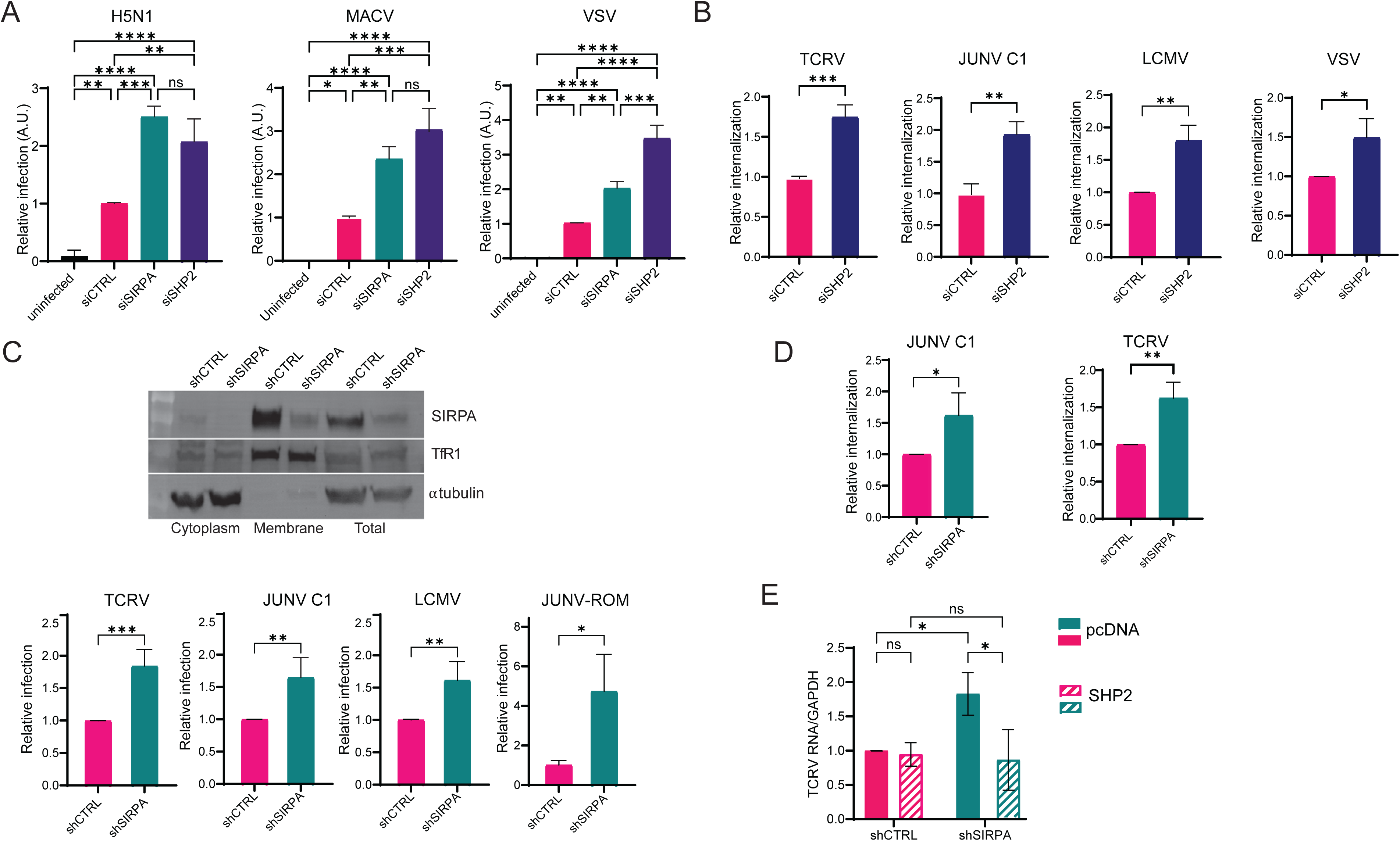
SHP2 is a virus restriction factor. A) A549 (H5N1) or U2OS (MACV, VSV) cells were transfected with the indicated siRNAs and then infected with H5N1, MACV, or VSV G pseudoviruses. Luciferase assays were performed at 48 hpi. Shown is the average ± SD of 3 independent experiments. One-way ANOVA was used to determine significance. ns, not significant, *, *P* ≤ 0.05; **, *P* ≤ 0.01; ***, *P* ≤ 0.001; ****, *P* ≤ 0.0001. B) VIAs were carried out with U2OS cells transfected with the indicated siRNAs. Internalized viral RNA was analyzed by RT-qPCR. Shown is the average ± SD of 3 independent experiments. Student’s T test was used to determine significance. *, *P* ≤ 0.05; **, *P* ≤ 0.01; ***, *P* ≤ 0.001. C) Top: SIRPA expression in different cellular fractions from shSIRPA or shcontrol (shCTRL) stably transduced U2OS cells was analyzed by western blot. Bottom: shSIRPA or shCTRL cells were infected with the indicated viruses. RNA was isolated at 24 hpi and viral nucleic acids were measured by RT-qPCR. Shown is the average ± (SD) of 3–4 independent experiments. Student’s T test was used to determine significance. *, *P* ≤ 0.05; **, *P* ≤ 0.01; ***, *P* ≤ 0.001. D) VIAs were carried out with shSIRPA or shCTRL cells. Internalized viral RNA was analyzed by RT-qPCR. Shown is the average ± SD of 3 independent experiments. Student’s T test was used to determine significance. *, *P* ≤ 0.05; **, *P* ≤ 0.01. E): shSIRPA or shCTRL cells were transfected with pcDNA or SH2 expression plasmids and infected with TCRV. RNA was isolated at 24 hpi and viral nucleic acids were measured by RT-qPCR. Shown is the average ± SD of 3 independent experiments. One-way ANOVA was used to determine significance. *, *P* ≤ 0.05.

We found previously that SIRPA blocked virus endocytosis (Sarute et al., 2021). To determine if SHP2, like SIRPA, played a role in restricting this step of infection, we performed virus internalization assays (VIA). Infectious TCRV, JUNV C1, LCMV and VSV were incubated with SHP2-depleted cells and at 1 hr post-binding, internalized virus RNA levels were measured. SHP2 depletion resulted in increased internalization of all 4 viruses (Fig. 1B). When we tested knockdown effects at later times in infection with replication-competent viruses, we found that SHP2 knockdown decreased virus levels, likely affecting a post-entry step in virus replication (not shown).

We next created stable U2OS cells expressing a SIRPA shRNA. Total and membrane associated SIRPA levels were depleted by shRNA knockdown (Fig. 1C). SIRPA knockdown had no effect on expression of the JUNV entry receptor TfR1 (Fig. 1C). At 24 hr post-infection TCRV, JUNV C1, LCMV and *bona fide* JUNV (Romero strain) levels were higher in the knockdown cells than in cells expressing control shRNA (Fig. 1C). Moreover, both JUNV C1 and TCRV internalization increased in the SIRPA knockdown cells (Fig. 1D). We then transfected these cells with a SHP2 expression vector. SHP2 overexpression in the SIRPA knockdown cells decreased TCRV infection levels back to those seen with shControl, as would be expected if SHP2 is downstream of SIRPA (Fig. 1E). Interestingly, SHP2 overexpression in SIRPA-expressing cells had no effect on TCRV infection, suggesting that SHP2 association with endogenous SIRPA was already maximal (Fig. 1E). Thus, like SIRPA, SHP2 plays a role in blocking endocytosis of viruses that traffic to an acidic compartment.

### SIRPA and SHP2 interact during virus infection

We then used co-immunoprecipitation and proximity ligation assays (PLA) to examine the interaction between SIRPA and SHP2 during infection. We co-transfected flag-tagged wild type SIRPA, SIRPA lacking the cytoplasmic tail (ΔCyt) or SIRPA with Y-to-A mutations in the ITIM and HA-tagged SHP2 expression constructs into U2OS cells and carried out co-immunoprecipitations; the Y-to-A ITIM and ΔCyt mutants were unable to restrict virus infection (Sarute et al., 2021). All SIRPA constructs except ΔCyt and Y-to-A ITIM interacted with SHP2 (Fig. 2A). This was confirmed by proximity ligation assays (PLA); while WT SIRPA interacted with SHP2, deletion of the cytoplasmic tail abrogated this interaction and the Y-to-A ITIM mutant showed significantly less interaction (Fig. S1A).

**Figure 2.**
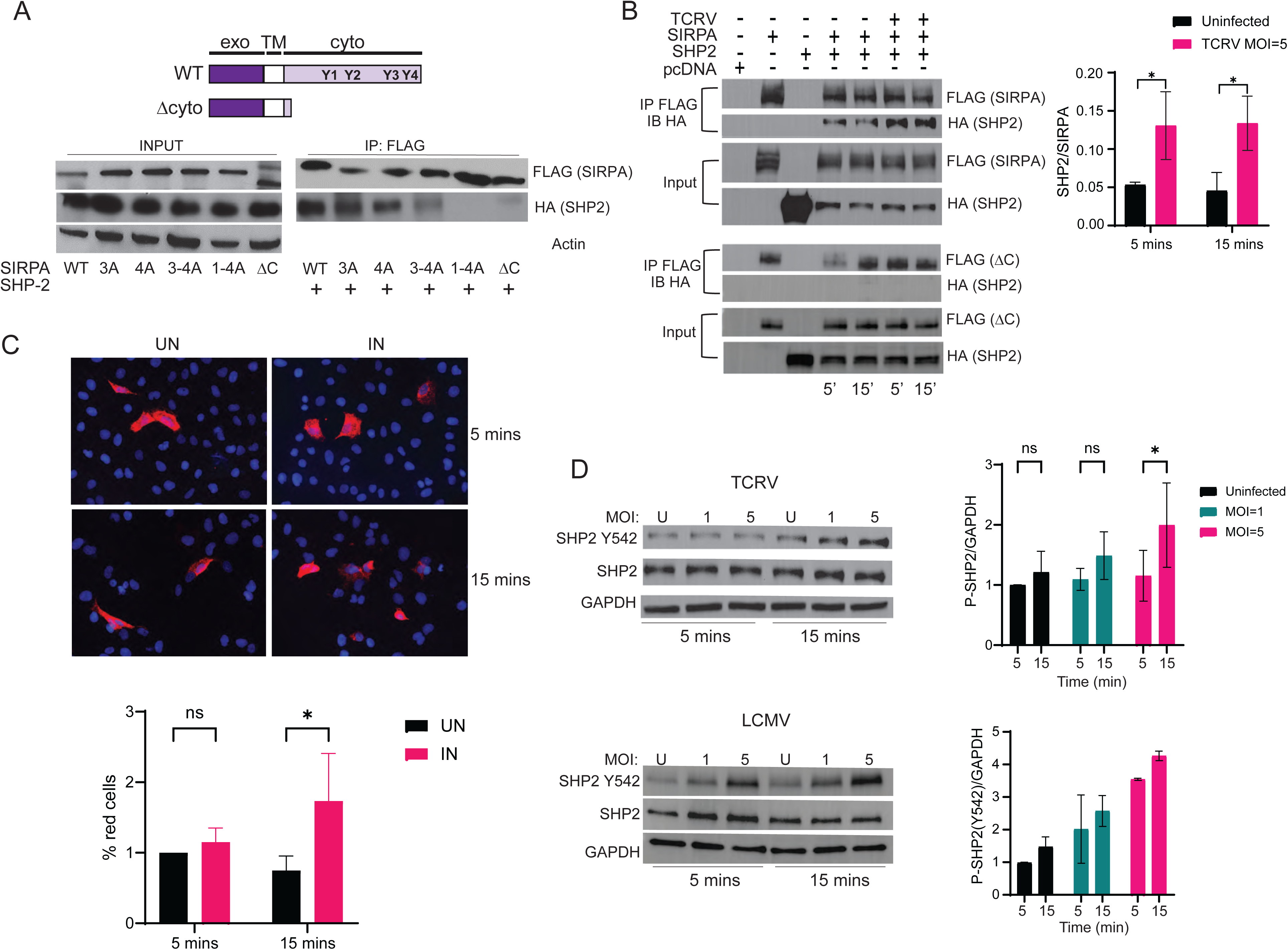
SHP2 is a virus restriction factor. A) U2OS cells were co-transfected with indicated Flag-tagged SIRPA and HA-tagged SHP2 expression plasmids. Cell lysates were immunoprecipitated with anti-Flag antibody, and western blots were subjected to probing with anti-HA antibody. B) U2OS cells transfected with SIRPA-FLAG or SHP2-HA constructs were infected with TCRV (MOI=5) for 5 or 15 minutes. Cell lysates were immunoprecipitated with anti-Flag antibody, and western blots were probed with anti-HA antibody. Quantification of SHP2 level normalized to SIRPA levels is shown in the bar graph (average ± SD of 3 independent experiments). One-way ANOVA was used to determine significance. *, *P* ≤ 0.05. C) U2OS cells transfected with SIRPA-FLAG or SHP2-HA were infected with TCRV (MOI=5) for 5 or 15 minutes. PLA using anti-Flag antibody and anti-HA antibody was carried out. Shown below the photographs is quantification of the appearance of the PLA^+^ spots (average ± SD of 3 independent experiments). For each experimental condition, at least eight fields were imaged. One-way ANOVA was used to determine significance. ns, not significant, *, *P* ≤ 0.05. D) U2OS cells were infected with TCRV or LCMV (MOI=1) or 5 for 5 or 15 minutes. Western blots were probed p-SHP2 (Y542) antibody. Quantification of phospho-SHP2 normalized to GAPDH (average ± SD of 3 [TCRV] or 2 [LCMV] independent experiments) is shown to the right. One-way ANOVA was used to determine significance. *, *P* ≤ 0.05.

To determine whether virus affected SIRPA-SHP2 interaction, SIRPA and SHP2-transfected U2OS cells were incubated with TCRV for 5 or 15 minutes, SIRPA was immunoprecipitated and western blots were performed. Although over-expressed SIRPA and SHP2 interacted in the absence of virus, more SHP2 co-immunoprecipitated with SIRPA after virus binding, at both 5– and 15-min post-incubation (Fig. 2B, top). No interaction with ΔCyt was seen after virus incubation (Fig. 2B, bottom). This interaction was also examined by PLA. Co-localization of the transfected proteins could be detected in the presence and absence of virus (Fig. 2C). Moreover, the co-localization signal was increased after 15-min incubation with TCRV (Fig. 2C).

SHP2 phosphorylation at tyrosine 542 is associated with its ability to interact with ITIM-containing proteins. We next tested if incubation of U2OS cells with virus altered the phosphorylation status of endogenous SHP2 and found that both TCRV and LCMV caused increased phosphorylation at Y542, especially by 15 min post-incubation (Fig. 2D).

Taken together, these data suggest that SIRPA and SHP2 interact during virus entry, activating SHP2 by phosphorylation and initiating a process that leads to down-regulation of virus endocytosis.

### SIRPA is phosphorylated during virus infection

Activation of SIRPA and recruitment of SHPs requires phosphorylation of its ITIMs (Tsuda et al., 1998). We next examined SIRPA’s phosphorylation state in the presence and absence of virus. All extracts examined for phospho-SIRPA were treated with deglycosylase prior to gel electrophoresis; SIRPA is a glycoprotein and the glycosylated, phosphorylated endogenous protein migrated as a diffuse, high molecular weight band that was difficult to quantify (Fig. S1B). U2OS cells were incubated for 5 and 15 min in the presence of TCRV or LCMV, the extracts were deglycosylated, endogenous SIRPA was immunoprecipitated and western blots were probed with anti-phospho-tyrosine antibody. While total SIRPA levels were unchanged, phospho-SIRPA levels doubled at 15 min. after incubation with either virus (Fig. 3A). Similar results were obtained with the mouse macrophage line NR9456 after incubation with JUNV C1 (Fig. S1C).

**Figure 3.**
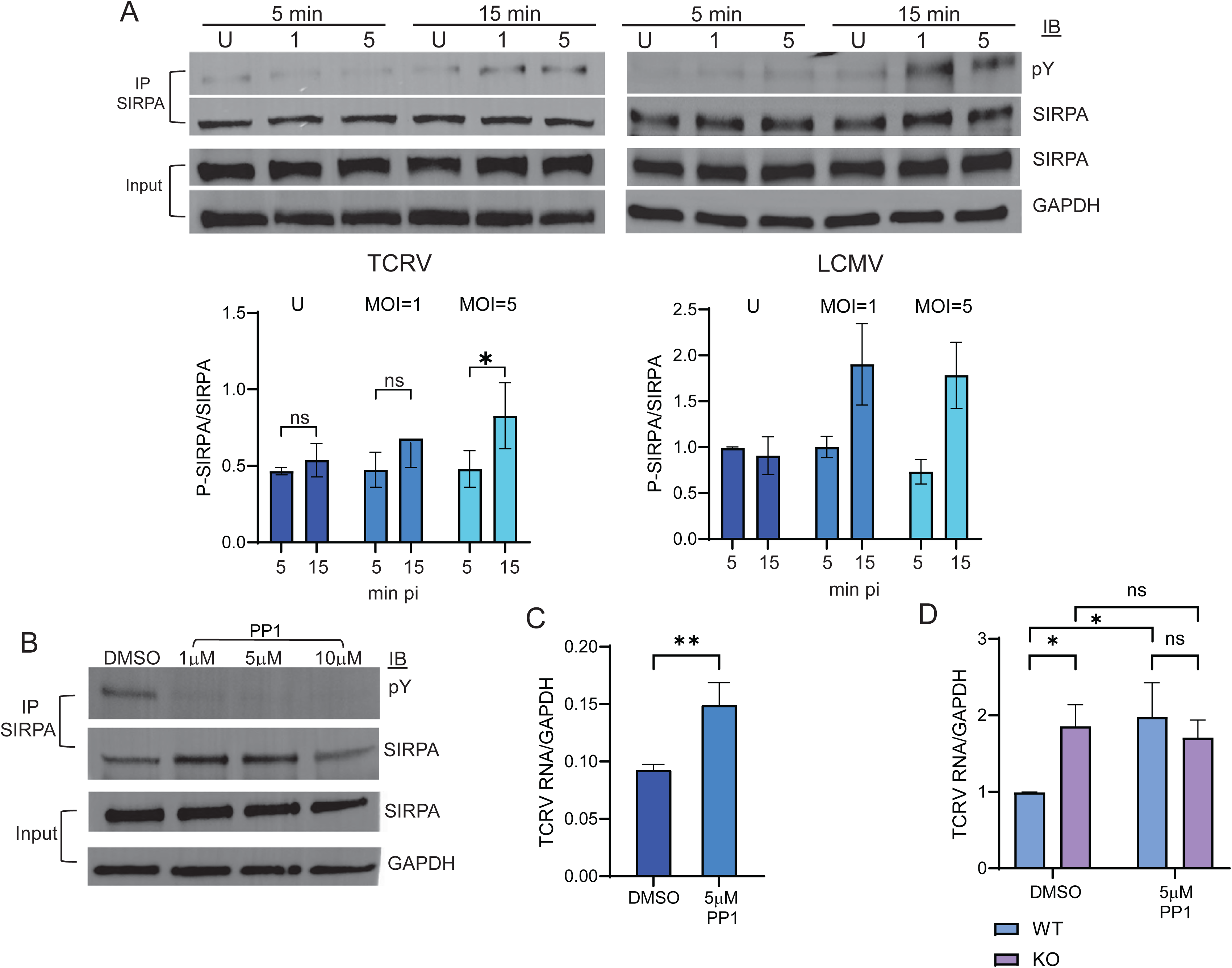
SIRPA is phosphorylated during virus infection. A) U2OS cells were infected with TCRV or LCMV (1 or 5 MOI, as indicated) for 5 or 15 minutes. Cell lysates were de-glycosylated cell lysates, immunoprecipitated with anti-SIRPA antibody, and western blots were probed with anti-phospho-tyrosine antibody. Quantification of phospho-SIRPA normalized to SIRPA (average ± SD of 3 [TCRV] or 2 [LCMV] independent experiments) are shown below. One-way ANOVA was used to determine significance. *, *P* ≤ 0.05. U, uninfected. B) U2OS cells were treated with PP1 inhibitor at the indicated concentrations. De-glycosylated cell lysates were immunoprecipitated with anti-SIRPA antibody, and western blots were subjected to probing with anti-phospho-tyrosine antibody. C) U2OS cells were pre-treated with PP1 inhibitor at the indicated concentrations for 1 hour and then infected with TCRV (MOI = 1) in the presence of inhibitor for 1 hour and virus and drug were removed. Viral nucleic acids were measured by RT-qPCR at 24 hpi. Shown is the average ± SD of 3 independent experiments. Student’s T test was used to determine significance. **, *P* ≤ 0.01. D) Fibroblasts isolated from mice of the indicated genotype were pre-treated with PP1 inhibitor at indicated concentration for 1 hour. Pretreated fibroblasts were then infected with TCRV (MOI=1) + PP1 inhibitor for 1 hour and incubated in fresh media for 24 hours. RNA was isolated at 24 hpi and examined by RT-qPCR. Shown is the average ± SD of 3 independent experiments. One-way ANOVA was used to determine significance. *, *P* ≤ 0.05.

SIRPA is phosphorylated by src family kinases (SFKs) (Tsuda et al., 1998). To determine if SFKs were responsible for the increase in SIRPA phosphorylation in response to virus, we tested if the pan-SFK inhibitor PP1 altered virus infection. First, we showed that PP1 treatment reduced basal phospho-SIRPA levels (Fig. 3B) as well as virus-induced phosphorylation (Fig. S1C). Moreover, PP1 treatment increased TCRV infection in U2OS cells (Fig. 3C). To ensure that PP1 was acting through SIRPA, we also tested fibroblasts derived from wild type and SIRPA knockout mice. PP1 only increased infection in WT and not knockout cells, which as we showed previously, are more highly infected by arenaviruses (Fig. 3D) (Sarute et al., 2021).

### FYN is required for SIRPA’s antiviral activity

We next determined which SFK might play a role in activating SIRPA’s antiviral activity. We used siRNA knockdown of *SRC*, *LYN*, *FYN* and *YES*, 4 ubiquitously expressed SFKs and tested the effect on infection by TCRV and JUNV C1. Depletion of *FYN* but not the other SFKs resulted in 2-fold increased infection by both viruses in U2OS cells, similar to what is seen with SIRPA knockdown (Fig. 4A). To ensure that *FYN* knockdown was affecting virus internalization, we also carried out VIA assays in knockdown cells. Knockdown of *FYN* increased TCRV and JUNV C1 internalization to levels similar to that seen with SIRPA or SH2 knockdown (Fig. 4B).

**Figure 4.**
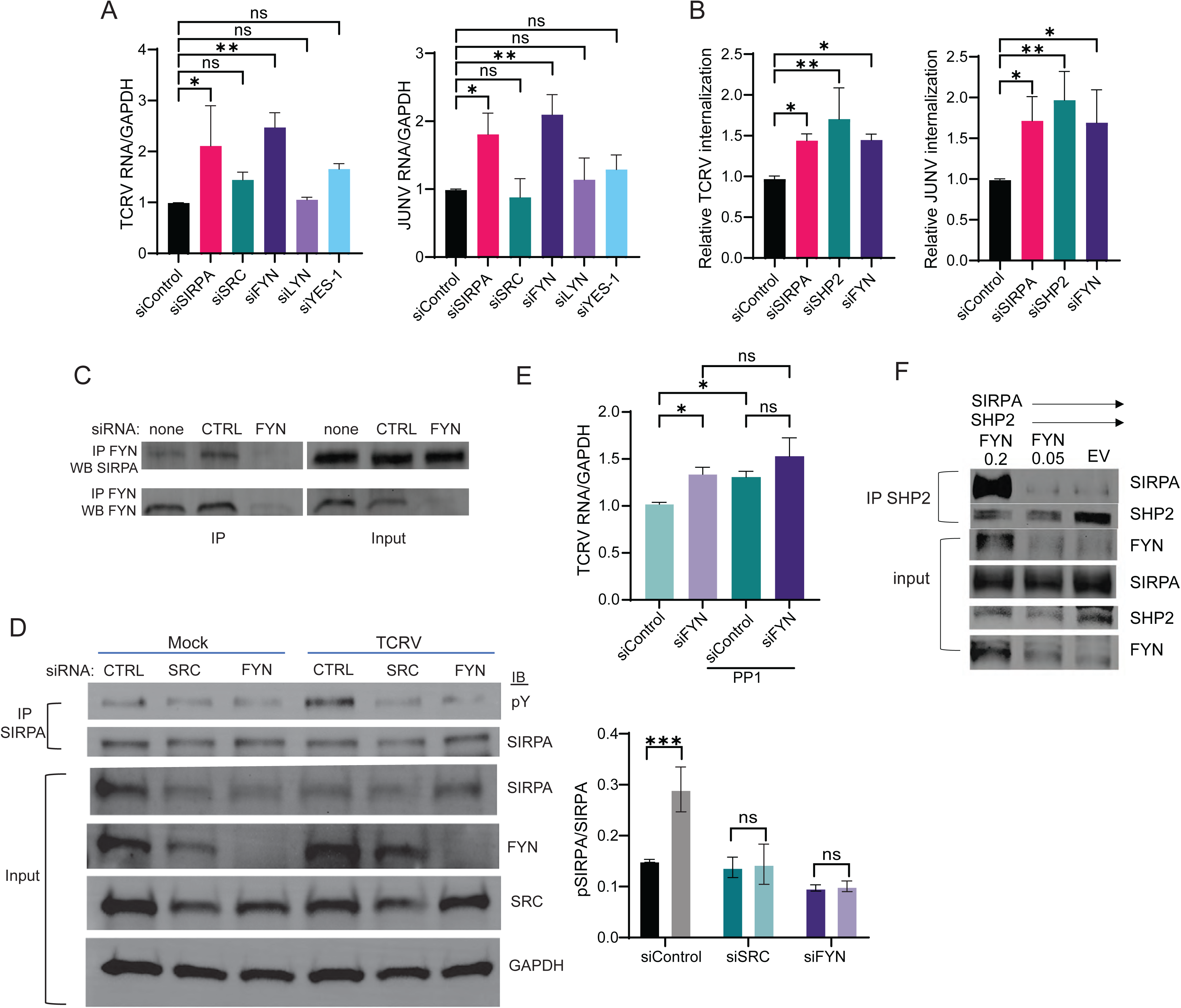
FYN is required for SIRPA’s antiviral activity. A) U2OS transfected with indicated siRNAs were infected with JUNV C1 or TCRV (MOI=1). RNA was isolated at 24 hpi and analyzed by RT-qPCR. Shown is the average ± SD of 3 independent experiments. One-way ANOVA was used to determine significance. *, *P* ≤ 0.05; **, *P* ≤ 0.01. B) VIAs were carried out with U2OS cells transfected with the indicated siRNAs and internalized JUNV C1 or TCRV RNA was analyzed by RT-qPCR. Shown is the average ± SD of 4 independent experiments. One-way ANOVA was used to determine significance. *, *P* ≤ 0.05; **, *P* ≤ 0.01. C) U2OS cells were transfected with the indicated siRNAs and immunoprecipitated and followed by western blotting with the indicated antibodies. D) U2OS cells transfected with indicated siRNA were infected with TCRV (MOI=1) for 5 for 5 or 15 minutes. De-glycosylated cell lysates were immunoprecipitated with anti-SIRPA antibody, and western blots were probed with anti-phospho-tyrosine antibody. Quantification of phospho-SIRPA normalized to total SIRPA is shown below (average ± SD of 3 independent experiments). One-way ANOVA was used to determine significance. ***, *P* ≤ 0.01. E) U2OS transfected with the indicated siRNAs were treated with 5 μM PP1 and infected with TCRV. Viral nucleic acids were measured by RT-qPCR at 24 hpi. One-way ANOVA was used to determine significance. *, *P* ≤ 0.05. F) U2OS cells were co-transfected with FYN, SIRPA and SHP2 expression vectors as indicated and co-IPs with anti-SHP2 antibody were performed.

Previous studies have suggested that SRC and FYN phosphorylate SIRPA (Tsuda et al., 1998). Over-expressed SRC, FYN and LYN all co-immunoprecipitated with SIRPA (Fig. S2A). This interaction depended on SIRPA’s ITIM, since neither the construct lacking the cytoplasmic tail or with 1-4A substitutions interacted with Fyn (Fig. S2B). Endogenous FYN and SIRPA also co-immunoprecipitated but not in extracts from cells which received FYN siRNAs (Fig. 4C).

We then tested if SRC or FYN altered SIRPA phosphorylation. Over-expression of SRC, FYN or LYN all increased SIRPA phosphorylation (Fig. S2C) and FYN knockdown decreased basal levels of phospho-SIRPA; SRC knockdown also decreased phosph-SIRPA, but to a lesser extent (Fig. S2D). Moreover, during virus infection, cells depleted for FYN or SRC showed lower levels of SIRPA phosphorylation after 15 min incubation with TCRV (Fig. 4D). Treatment of cells depleted for FYN with PPI did not further increase TCRV infection (Fig. 4E). Finally, over-expression of FYN increased the interaction between SIRPA and SHP2 in co-immunoprecipitation assays (Fig. 4F). Taken together, these data show that SFKs, especially FYN, are needed for SIRPA phosphorylation and recruitment of SHP2 during virus infection.

### The role of integrin in virus internalization

Upregulating integrin levels by treating cells with MnCl_2_ increased infection by JUNV C1 and LCMV (Sarute et al., 2021). We next tested the integrin inhibitor BTT-3033. BTT-3033 concentrations up to 10μM had no effect on cell viability (Fig. S3A). BTT-3033 decreased infection by TCRV, LCMV and JUNV C1 in U2OS cells (Fig. 5A). We also tested whether BTT-3033 could JUNV-Romero infection and found that it was as effective as the fusion inhibitor LHF-535, which inhibits TCRV and LASV and which we previously showed was effective in treating JUNV-Romero infection in guinea pigs (Madu et al., 2018; Westover et al., 2024) (Fig. S3A). This inhibition was not limited to arenaviruses, as ZIKA virus (ZIKV) infection was also diminished by BTT-3033 treatment (Fig. S3B); other integrin inhibitors have previously been reported to inhibit ZIKV infection (Wang et al., 2020). In addition to testing the BTT-3033 effects in wild type cells, we used SIRPA shRNA or siRNA knockdown cells. BT-3033 decreased infection by TCRV, JUNV C1 and LCMV in SIRPA shRNA knockdown cells, reducing it to levels similar to that seen in wild type cells (Fig. 5A). BTT-3033 also inhibited TCRV and JUNV C1 internalization (Fig. 5B) and reduced infectious JUNV C1 titers produced by both shCTRL and shSIRPA knockdown cells (Fig. 5C). MACV GP and VSV G pseudovirus infection was also inhibited by BT-3033 in both siControl and siSIRPA treated cells (Fig. 5D).

**Figure 5.**
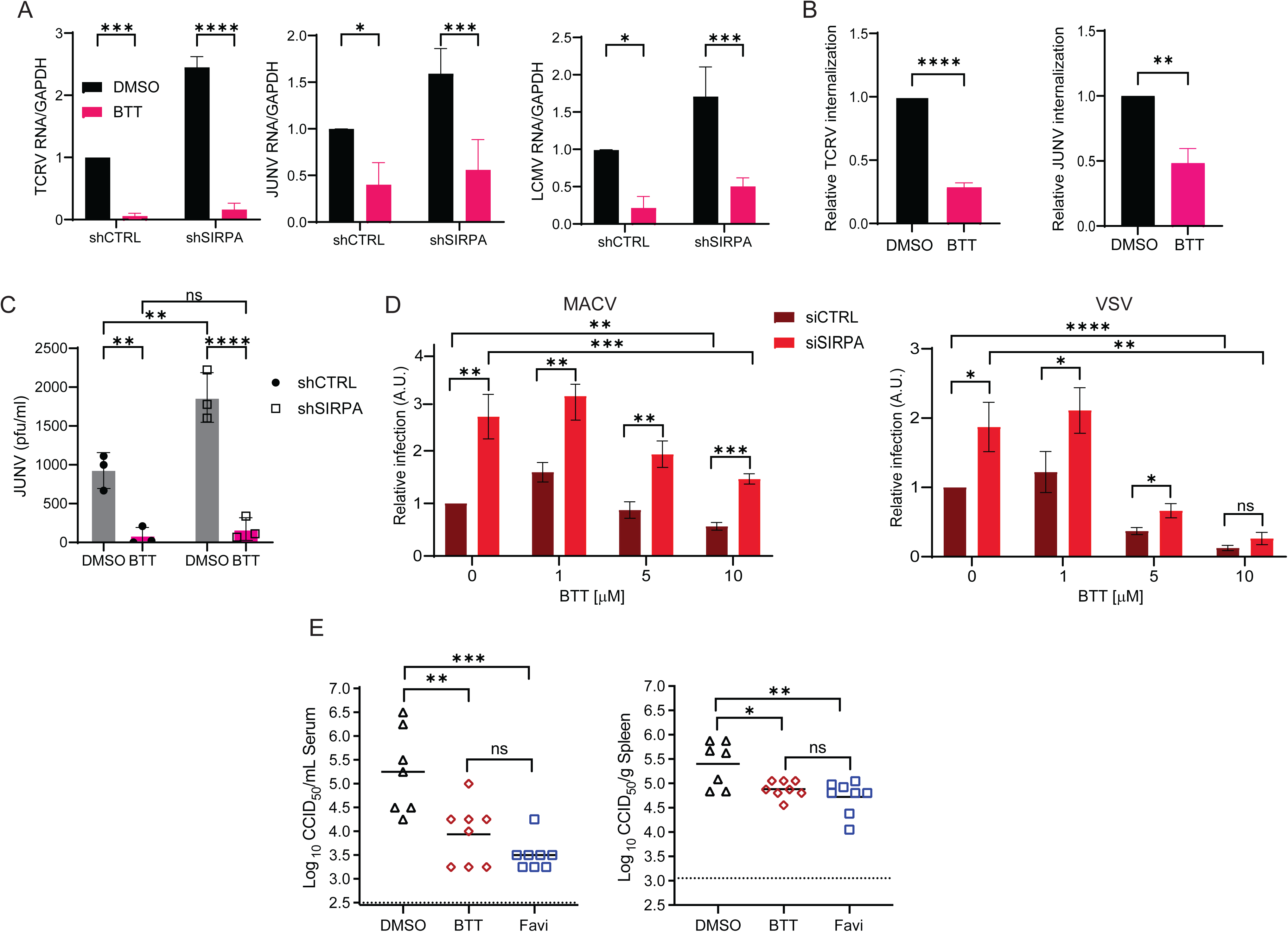
The integrin pathway and virus internalization. A) shSIRPA or shCTRL cells were treated with BTT-3033 and infected with the indicated viruses. RNA was isolated at 24 hpi and viral RNA was measured by RT-qPCR. Shown is the average ± SD of 3–4 independent experiments. One-way ANOVA was used to determine significance. *, *P* ≤ 0.05; ***, *P* ≤ 0.001; ****, *P* ≤ 0.0001. B) VIAs were carried out with U2OS cells treated with BTT-3033. Internalized viral RNA was analyzed by RT-qPCR. Shown is the average ± SD of 3 independent experiments. Student’s T test was used to determine significance. **, *P* ≤ 0.01; ****, *P* ≤ 0.0001. C) shSIRPA or shCTRL cells were treated with BTT-3033 and infected with JUNV C1. Viral titers were analyzed at 24 hpi. Shown is the average ± SD of 3 independent experiments. One-way ANOVA was used to determine significance. ns, not significant, **, *P* ≤ 0.01; ****, *P* ≤ 0.0001. D) U2OS cells were treated with BTT-3033 at indicated concentrations and infected with MACV or VSV pseudoviruses. Luciferase assays were performed at 48 hpi. Shown is the average ± SD of 3 independent experiments. One-way ANOVA was used to determine significance. *, *P* ≤ 0.05; **, *P* ≤ 0.01; ***, *P* ≤ 0.001; ****, *P* ≤ 0.0001. E) Viral titers in serum and spleen of BTT-3033– and favipravir-treated mice. Horizontal lines represent the mean for each group. The dotted horizontal lines represent the assay’s lower limit of detection. \**, P* ≤ 0.05; \*\**, P* ≤ 0.01; \*\*\**, P* ≤ 0.001 compared between groups. One animal in the DMSO placebo group was excluded from the analysis due to complications unrelated to the viral infection.

We also tested if BTT-3033 would diminish *in vivo* JUNV C1 infection. Wild type mice rapidly clear JUNV C1 infection, so we used transgenic mice expressing the human TfR1 (huTfR1 Tg mice), which are more susceptible to infection with JUNV (Hickerson et al., 2022). In addition, the mice were pre-treated with anti-interferon type 1 receptor antibody 24 hr prior to inoculation with virus to further enhance infection (Hickerson et al., 2020). Groups of four-week-old mice received once-daily intraperitoneal (IP) injections of BTT-3033 (1 mg/kg body weight) or DMSO vehicle 1 hour prior to challenge with approximately 10^5^ CCID_50_ (median cell culture infectious dose) of JUNV C1. As a control, a group of mice received favipiravir (100 mg/kg), twice daily, starting 1 hour prior to virus challenge; favipiravir is a known inhibitor of NWA infection *in vivo* (Gowen et al., 2013; Westover et al., 2024). The groups of mice were treated daily until 6-days post-infection, at which time serum and spleen were collected to assess viral loads. BTT-3-33 reduced JUNV Cl1 titers in both spleen and sera to the same level as favipiravir (Fig. 5E).

These data support our hypothesis that SIRPA works through blocking integrin-mediated virus endocytosis.

### FAK inhibition decreases virus infection

Integrin signaling works through phosphorylating FAK in many cell types, leading to downstream signaling and generation of the phagocytic cup (Larsen et al., 2003; Morrissey et al., 2020). We first tested whether virus interaction with cells altered FAK phosphorylation. U2OS cells were incubated with TCRV for 5 or 15 min and western blots were performed, using an antibody against pFAK Y397, an activating autophosphorylation required for downstream signaling (Golubovskaya et al., 2008). FAK phosphorylation was increased by incubation with either 1 or 5 MOI virus (Fig. 6A).

**Figure 6.**
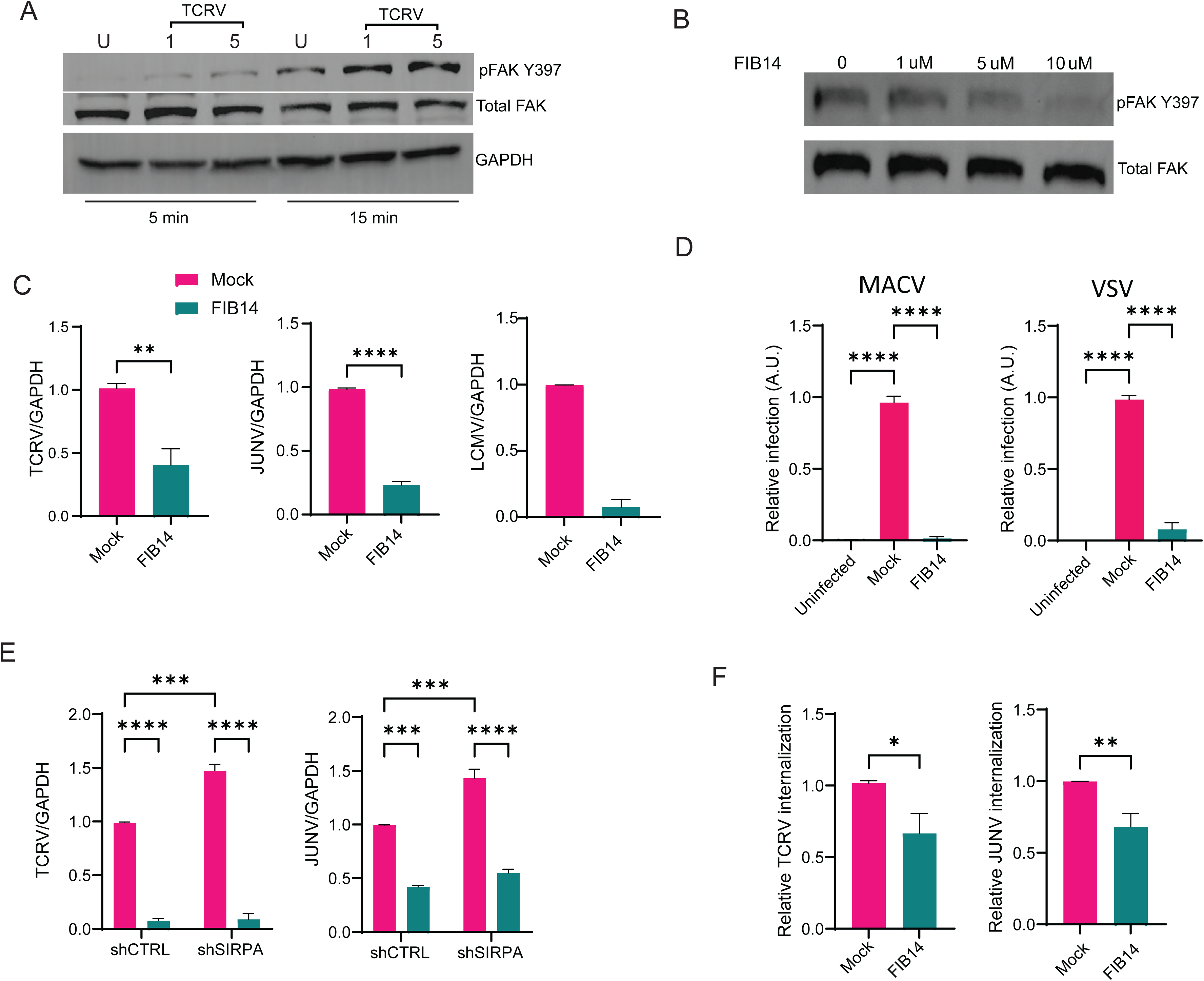
FAK signaling in virus internalization. A) U2OS cells were infected with TCRV (MOI=1) for 5 or 15 minutes. Western blots were probed with phospho-FAK (Y397) antibody. U, uninfected. B) U2OS cells were treated with FIB-14 for 1 hour and western blots were probed phospho-FAK (Y397) antibody. C) U2OS were treated with FIB-14 and infected with the indicated viruses. RNA was isolated at 24 hpi. Shown is the average ± SD of 3 (TCRV, JUNV C1) or 2 (LCMV) independent experiments. Student’s T test was used to determine significance. **, *P* ≤ 0.01; ****, *P* ≤ 0.0001. D) U2OS cells were treated with FIB-14 and infected with MACV, or VSV G pseudoviruses. Luciferase assays were performed at 48 hpi. Shown is the average ± SD of 3 independent experiments. One-way ANOVA was used to determine significance. ****, *P* ≤ 0.0001. E) shSIRPA or shCTRL cells were treated with FIB-14 and infected with the indicated viruses. RNA was isolated at 24 hpi. Shown is the average ± SD of 3 independent experiments. One-way ANOVA was used to determine significance. ***, *P* ≤ 0.001; ****, *P* ≤ 0.0001. F) VIAs were carried out with U2OS cells treated with FIB-14. Internalized viral RNA was analyzed by RT-qPCR. Shown is the average ± SD of 3 independent experiments. Student’s T test was used to determine significance. *, *P* ≤ 0.05; **, *P* ≤ 0.01.

If integrin signaling through FAK was also required for virus endocytosis, then loss or inhibition of FAK should result in decreased infection. We first tested whether knockdown of FAK altered infection by JUNV C1, LCMV or VSV. Surprisingly, cells treated with FAK siRNAs had higher levels of infection (Fig. S4A). However, when we looked at SIRPA RNA levels, we found that depletion of FAK resulted in decreased SIRPA expression, as did depletion of integrin alpha 5 (ITGA5) (Fig. S4B). Although we do not yet know mechanistically why this occurs, it confirms that loss of SIRPA leads to increased infection.

To bypass the problem of FAK depletion, we turned to the inhibitor FIB-14, which blocks activating FAK autophosphorylation on residue Y397 (Fig. 6B) (Golubovskaya et al., 2008). First, we tested whether blocking FAK phosphorylation altered SIRPA expression. Whereas FIB-14 concentrations of FIB-14 > 5 μM increased SIRPA protein levels at the cell surface, treatment of cells for 3 hr with 1 μM FIB-14 had minimal significant effect on surface expression (Fig. S4C). We incubated cells for 1 hr with 1 μM FIB-14, followed by 1 hr infection with TCRV, JUNV C1 or LCMV in the presence of drug. The cells were then washed to remove drug and virus, and infection was allowed to proceed for 24 hr. Examination of virus RNA levels demonstrated that all three viruses were inhibited by FIB-14 (Fig. 6C). FIB-14 also inhibited infection by MACV GP and VSV G pseudotypes (Fig. 6D). Moreover, FIB-14 treatment decreased TCRV and JUNV C1 infection in SIRPA knockdown cells (Fig. 6E). We also tested FIB-14’s effect on JUNV C1 and TCRV internalization and showed that both were reduced by drug treatment (Fig. 6F). Thus, FAK activation is needed for virus endocytosis and entry.

### Role of integrins in SIRPA anti-viral activity in macrophages

Sentinel cells of the immune system, such as macrophages, are likely early targets of infection by many RNA viruses *in vivo*, including new world arenaviruses (Cuevas and Ross, 2014; Martinez et al., 2012; Sarute and Ross, 2017). SIRPA restricts infection of immortalized and primary macrophages by New and Old World arenaviruses, VSV and ZIKV (Sarute et al., 2021).

To examine the overlap between the downstream pathway utilized by SIRPA in U2OS cells and macrophages, we used the immortalized monocytic cell line THP1, which can be differentiated to M1 and M2 macrophages with interferon γ (IFNγ) or interleukin 4 (IL-4), respectively (Spencer et al., 2010). We tested whether macrophage differentiation altered SIRPA expression and thus JUNV C1 infection. Differentiation to either M1 or M2 macrophages resulted in increased SIRPA surface expression and decreased JUNV C1 internalization (Fig. 7A).

**Fig. 7.**
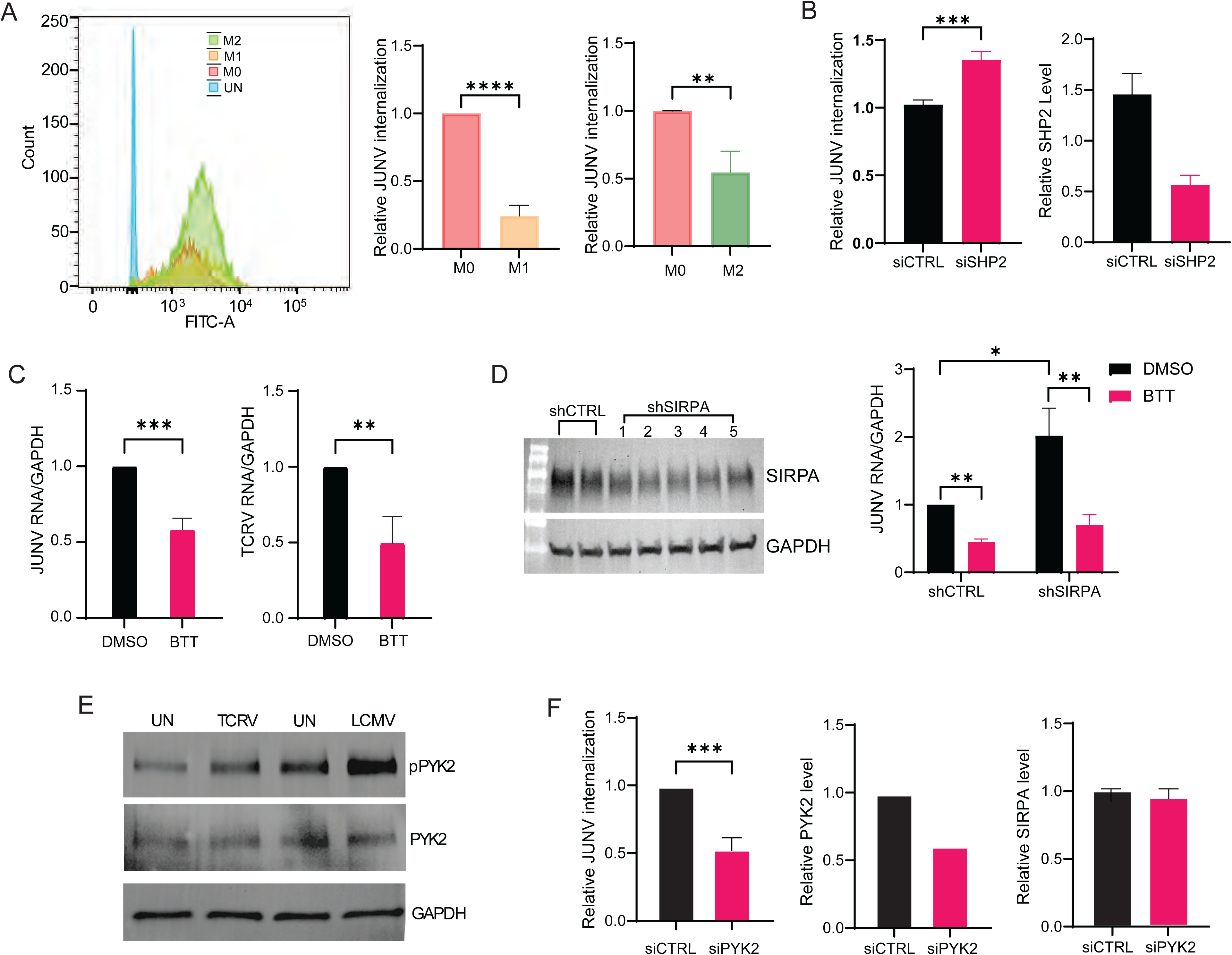
The integrin pathway is also important for infection of THP1 macrophages. A) Differentiation of THP1 macrophages to M1 or M2 phenotypes increase SIRPA expression and decreases JUNV C1 infection. Student’s T test was used to determine significance. ***, *P* ≤ 0.05; **, *P* ≤ 0.002. B) Knockdown of SHP2 in M0 THP1 cells increases infection. Student’s T test with Welch’s correction was used to determine significance. ***, *P* ≤ 0.0005. C) BTT-3033 decreases JUNV C1 and TCRV infection in M0 THP1 cells. Student’s T test with Welch’s correction was used to determine significance. **, *P* ≤ 0.007; ***, *P* ≤ 0.0007. D) BTT-3033 decreases infection in M2 THP1 cells SIRPA knockdown cells. Left – western blots showing SIRPA expression in 5 different clones. Clone 5 was used for the infection experiment in the right panel. SIRPA surface expression for #5 is shown in Suppl. Fig. 6B. Two-way ANOVA was used to determine significance. *, *P* ≤ 0.02; **, *P* ≤ 0.002. E) PYK2 is more highly phosphorylated after incubation with TCRV or LCMV for 15 min. F) Knockdown of PYK2 in M0 THP1 cells causes lower levels of JUNV C1 internalization. Student’s T test was used to determine significance. ***, *P* ≤ 0.007.

Next, we tested whether SHPs played a role in SIRPA’s inhibition of infection in macrophages. We were unable to test SHP1, because knockdown of this protein in THP1 cells caused growth arrest (not shown). We used M0 cells, which expressed the lowest levels of SIRPA. SHP2 knockdown had a modest but significant effect on JUNV internalization in THP1 M0 cells (Fig. 7B).

We also tested if the integrin inhibitor BTT-3033 inhibited JUNV C1 or TCRV infection of THP1 cells, first using M0 macrophages. JUNV C1 and TCRV infection was inhibited in THP1 M0 cells (Fig. 7C). We also depleted SIRPA in THP1 cells using shRNA and showed that even about a 50% decreased in SIRPA (Fig. 7D, left panel; Fig. S5A) increased JUNV C1 infection by 2-fold in M2 macrophages (Fig. 7D, right panel). Similar to what was seen with U2OS cells, BTT-3033 diminished infection independent of SIRPA levels in M2 macrophages (Fig. 7D, right panel). BTT-3033 also inhibited THP1 M2 cells’ ability to phagocytose beads, demonstrating that integrin signaling was essential to both processes (Fig. S5B). As expected, SIRPA knockdown did not alter bead phagocytosis, since CD47/SIRPA binding is required to inhibit this process (Fig. S5C).

We also tested if focal adhesion kinases played a role in virus internalization. Protein tyrosine kinase 2 (PYK2) is believed to be the primary downstream modulator of integrin signaling in macrophages and has been shown to associate with SIRPA in this cell type (Duong and Rodan, 2000; Ostergaard and Lysechko, 2005; Timms et al., 1999). Like FAK, activation of PYK2 results in phosphorylation of a critical tyrosine residue, Y402 (Butler and Blystone, 2005). First, we showed that incubation of cells with either TCRV or LCMV caused phosphorylation of Y402 in M0 THP1 cells (Fig. 7E). Moreover, knockdown of PYK2 in THP1 cells decreased JUNV C1 internalization, without affecting SIRPA levels (Fig. 7F). Taken together, these data suggest that virus internalization is dependent on the integrin pathway, and that SIRPA acts to inhibit this pathway in macrophages and other cell types.

## Discussion

Many enveloped viruses require trafficking to acidic compartments where membrane fusion between the virus and cell results in entry. These viruses require initial binding of their glycoproteins to cell surface receptors and then through largely unknown means, induce clathrin– or caveolin-mediated endocytosis, or micro– or macropinocytosis, resulting in the formation of virus-containing vesicles that traffic to the endosome/lysosome (Helenius, 2018). Similarly, phagocytosis is initiated by the binding of cell surface receptors to ligands on the surface of target particles or cells and results in trafficking of cargo-containing vesicles to acidic, degradative compartments (Boero et al., 2023; Fitzer-Attas et al., 2000; Vorselen, 2022). SIRPA, a potent inhibitor of phagocytosis, localizes to the phagocytic synapse upon activation, where it blocks integrin-mediated engulfment of targets (Morrissey et al., 2020). Since virus entry is also inhibited by SIRPA activation, this suggests that there is overlap at the early steps of both processes (Sarute et al., 2021; Sarute et al., 2019). Here we show that like phagocytosis, the integrin pathway plays a role in virus entry and that SIRPA likely blocks its signaling and thereby inhibits infection.

We tested this by systematically examining the role of downstream effectors of SIRPA-mediated inhibition of phagocytosis and integrin signaling in virus entry. Because many of the proteins that function in the integrin pathway likely affect multiple steps of virus replication, as we saw with SHP2 knockdown, we treated cells with inhibitors only during virus entry and then removed it from the media. We also validated the infection studies with VIAs, to ensure that the effects were on virus entry and not later stages of replication.

It is well-established that tyrosine phosphorylation of SIRPA’s ITIM by SFKs are required for the recruitment of SHPs to this domain, although which SFK is operative may depend on context and cell type (Barclay and Van den Berg, 2014; Kelley and Ravichandran, 2021). We found that SIRPA was phosphorylated upon binding of virus to cells and that PP1, a pan-SFK inhibitor inhibited SIRPA phosphorylation and increased virus entry. Several SFKs could phosphorylate SIRPA, particularly when over-expressed. Both FYN and SRC have been previously implicated in SIRPA phosphorylation (Tsuda et al., 1998). However, only FYN-deficient cells consistently had higher levels of infection than cells depleted for the other ubiquitously expressed SFKs SRC, YES and LYN.

The mechanism by which virus entry triggers SFK-dependent phosphorylation of SIRPA remains to be determined. Viruses like SARS-CoV-2 and Ebola, which are both inhibited by SIRPA, interact with cell surface integrins through their Arg-Gly-Asp (RGD) domains, which plays a role in their entry and could also activate SIRPA (Hussein et al., 2015; Liu et al., 2022; Schornberg et al., 2009). Moreover, previous studies have shown that binding to phosphatidyl serine (PS) on the surface of apoptotic cells transduces a signal leading to SIRPA phosphorylation (Tada et al., 2003). Many of the viruses inhibited by SIRPA are thought to incorporate PS in their membranes, which through binding to PS receptors may facilitate infection (Bohan and Maury, 2021; Jemielity et al., 2013; Shimojima et al., 2012). This may be one means by which virus binding to cells activates SIRPA. We showed previously that CD47 on virions does not trigger SIRPA-mediated inhibition of infection, but a recent study suggests that the arenavirus LCMV may be transmitted cell-to-cell, which could result in CD47 on infected cells activating SIRPA on uninfected cells and thereby suppressing virus transmission *in vivo* (Byford et al., 2024; Sarute et al., 2021)..

We previously showed that SIRPA’s ITIM was required for its inhibition of infection and here showed that it also was needed to bind SHP2 (Sarute et al., 2021). Given that both SIRPA and SHP2 restrict virus entry, we investigated whether SIRPA and SHP2 interact during viral entry. NW arenaviruses are internalized within 30 minutes of incubation with virus (Lavanya et al., 2013). We therefore selected early time points to examine this interaction and showed that SIRPA-SHP2 interaction increased within minutes of virus infection; this coincided with SHP2 tyrosine phosphorylation at Y542, which is associated with SHP2 activation. Both SIRPA phosphorylation levels and SIRPA-SHP2 interaction significantly decreased in FYN-deficient cells, supporting a model in which virus-activated phosphorylation of SIRPA initiates a downstream signaling pathway through recruiting SHPs (Fig. 7).

Phosphorylation of SIRPA’s ITIM region recruits and activates the non-receptor tyrosine phosphatases SHP1 and SHP2 to inhibit phagocytosis (Granda Farias et al., 2025; Li et al., 2023; Murata et al., 2014). While SHP1 is primarily expressed in hematopoietic cells, SHP2 is ubiquitously expressed (Lorenz, 2009). Our experiments, primarily conducted in the osteosarcoma cell line U2OS, focused on the SHP2 protein. We showed previously that SHP2 depletion led to decreased virus infection levels (Sarute et al., 2019). However, SHP proteins function as negative regulators in a wide range of signaling pathways and could affect steps in virus replication beyond entry. Since SIRPA blocks virus endocytosis we determined if SHP2 deficiency specifically affected virus entry. H5N1, MACV, and VSV pseudovirus all showed significantly increased infection and JUNV-C1, TCRV, LCMV, and VSV internalization increased in SHP2-deficient U2OS or A549 cells. Moreover, overexpression of SHP2 in SIRPA shRNA but not control cells decreased infection. These findings suggest that, like SIRPA, SHP2 decreases virus entry but also likely affects downstream steps in virus replication.

Activated SIRPA is localized to the phagocytic synapse where it prevents integrin activation and thereby inhibits phagocytosis (Morrissey et al., 2020). Activation of integrin by Mn^2+^ treatment reverses SIRPA’s inhibition of phagocytosis, and we also showed that Mn^2+^ overcame SIRPA’s antiviral activity (Morrissey et al., 2020; Sarute et al., 2021). Depletion of integrin or other components in the signaling pathway is likely to alter many cellular functions, making it difficult to determine the direct effects on virus entry. To bypass this, we used short-term treatment of cells with integrin and integrin pathway inhibitors and studied their effects on virus internalization and infection. Both the integrin inhibitor BT-3033, which has previously been shown to inhibit SARS-CoV-2 infection, and the FAK inhibitor FIB-14 decreased virus infection and internalization, regardless of the presence of SIRPA (Simons et al., 2021).

FAK is believed to be a critical component of integrin signaling and is a target of SHPs; phospho-FAK associates with integrin and SHP proteins are believed to dephosphorylate it and thereby abolish integrin mediated-functions like adhesion and migration and actin polymerization needed for cytoskeletal rearrangement and phagocytic synapse formation (Larsen et al., 2003). FAK has been implicated in influenza virus replication and egress, and although we did not test it directly here, likely also plays a role in its entry, given that this step is inhibited by SIRPA and SHP2 (Fig. 1) (Elbahesh et al., 2016; Elbahesh et al., 2014; Sarute et al., 2021).

As macrophages are believed to be the initial targets of infection *in vivo* by many RNA viruses, we also tested whether SIRPA, SHPs, integrin and the focal adhesion kinase PYK2 played a role in JUNV C1 infection of THP1 cells. We found that M1 or M2 macrophages derived from THP1 cells expressed higher levels of SIRPA and thus sustained lower levels of virus infection than M0 cells. Moreover, although we were unable to test the role of the major hematopoietic SHP protein SHP1 in these cells, knockdown of SHP2 did increase JUNV C1 internalization to a small extent. Moreover, both inhibition of integrin signaling by BT-3033 and knockdown of PYK2, a FAK family protein expressed in macrophages, decreased infection of these cells and incubation with virus increased PYK2 phosphorylation. These data suggest that SIRPA inhibition of integrin signaling is likely to occur in multiple cell types, including biologically relevant targets like macrophages.

Interestingly, it was recently shown that CD47 binding to SIRPA on macrophages slows phagocytosis at the engulfment step but not at the binding or initiation step, and that this was dependent on SHP phosphatases (Miller et al., 2025). Moreover, it was suggested that the GTP exchange factor Vav1 was the target of SIRPA-bound SHPs and that the signal initiated by CD47 binding inhibited Rac, a known activator of non-muscle myosin II activation (NMII), which has also been implicated in phagocytosis (Tsai and Discher, 2008). NMII activation is also required for entry of various RNA viruses, including NWA (Ansin et al., 2025).

In conclusion, we propose a model in which virus entry triggers SFK-dependent phosphorylation of SIRPA’s ITIM region. This phosphorylation recruits and activates SHP2, which then functions downstream to restrict integrin signaling and inhibit virus endocytosis (Fig. 8). This model suggests that several steps in this pathway are targets for therapeutic intervention.

**Fig. 8.**
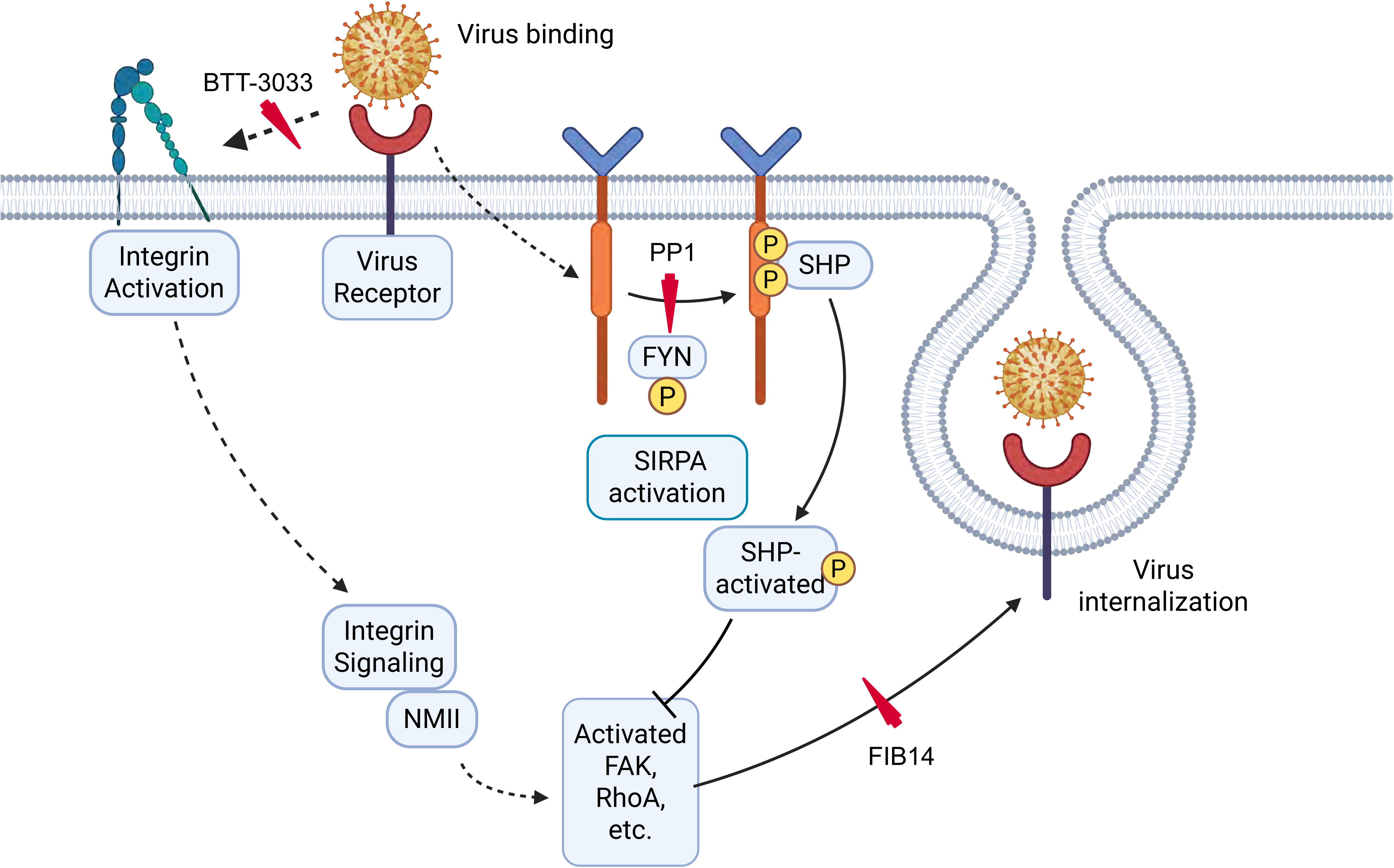
RNA viruses rely on integrin-mediated signaling to form the cup needed for virus endocytosis. This step is inhibited by integrin inhibitors such as BT-3033 and the FAK inhibitor FIB-14. When virus binds to cells, an unknown signal also triggers Fyn to phosphorylate tyrosine residues in SIRPA’s ITIM region. This step is inhibited by SRC family inhibitors like PP1. SIRPA phosphorylation recruits and activates SHP2, which in turn dephosphorylates signaling molecules in the integrin signaling pathway like FAK and NMII. The dephosphorylation of integrin pathway components negatively impacts cytoskeletal rearrangement and actin polymerization, thereby reducing viral internalization. Originally created in BioRender. Yan, M. (2024) BioRender.com/h75b068 and modified by Ross, S. (2026) https://BioRender.com/ ooyrvzl.

## Materials and methods

### Mice

B6.129P2 Sirpa<tm1Nog>/Rbrc (SIRPA KO) mice, lacking exons 7 and 8 in the *Sirpa* gene, were originally obtained from the Riken Institute (Japan) and bred to C56BL/6N mice at the University of Illinois (Inagaki et al., 2000). SIRPA wild type (WT) mice were originally derived by crossing mice heterozygous for the SIRPA KO allele, as previously described (Sarute et al., 2021). SIRPA KO and SIRPA WT mice were maintained as separate strains. For some experiments, C57BL/6N mice were used as WT controls. The mice were housed according to the policies of the Animal Care Committee of the University of Illinois at Chicago; all studies were performed in accordance with the recommendations in the Guide for the Care and Use of Laboratory Animals of the National Institutes of Health. The experiments performed with mice in this study were approved by University of Illinois at Chicago ACC protocol #24-111.

The huTfR1 Tg mice were obtained from Genentech (Yu et al., 2014), and the homozygous animals used in our studies have been previously described (Hickerson et al., 2020). The mice were bred at USU, and the genotype was confirmed by PCR. Animal procedures used in this study complied with guidelines set by the USDA and the Utah State University Institutional Animal Care and Use Committee.

### Cell lines

Vero E6, U2OS, A549, THP-1, and 293T cells were obtained from ATCC; NR-9456 cells were obtained from BEI Resources, NIAID/NIH. Vero E6, U2OS, A549, NR-9456 and 293T cells were cultured in Dulbecco’s modified Eagle Medium (DMEM; Gibco) supplemented with 2 mM glutamine, 10% fetal bovine serum (FBS, Invitrogen), and penicillin (100 U/ml)-streptomycin (100 μg/ml) (complete DMEM media). THP-1 cells (ATCC) were cultured in RPMI-1640 medium (ATCC 30-2001) with β-mercaptoethanol (0.05 mM) (BioRad) and 10% FBS (Invitrogen). The cells were incubated with 30ng/mL phorbol 12-myristate 13-acetate (PMA; Sigma Aldrich #P1585) for 24 hr to generate M0 macrophages, and then further differentiated to M1 macrophages using lipopolysaccharide (from *Escherichia coli* O55:B5; Millipore Sigma #L4524) and IFNγ (Millipore Sigma #l17001)(15 ng/ml and 50 ng/ml) or recombinant IL-4 (100 ng/ml; PeproTech/Thermofisher #200-04) for 48 hrs to generate M1 and M2 macrophages, respectively.

### RNA interference

The following siRNAs were used for gene depletion: human SIRPA (Ambion Silencer Select AM51331), mouse SIRPA (Dharmacon #D-042804-04-0002), siRNA control (Qiagen #1022076), human SHP2 (Qiagen #100044002), human FYN (Qiagen #102654729), human SRC (Qiagen #102223921), human LYN (Qiagen #100605577), human YES-1 (Qiagen #102223942). Cells were transfected at 60–70% confluence with the aforementioned siRNAs using the Lipofectamine RNAiMax (Invitrogen) forward transfection method for 48 hr. Knockdowns were confirmed by RT-qPCR with the primers described in Table 1.

**Table 1.**
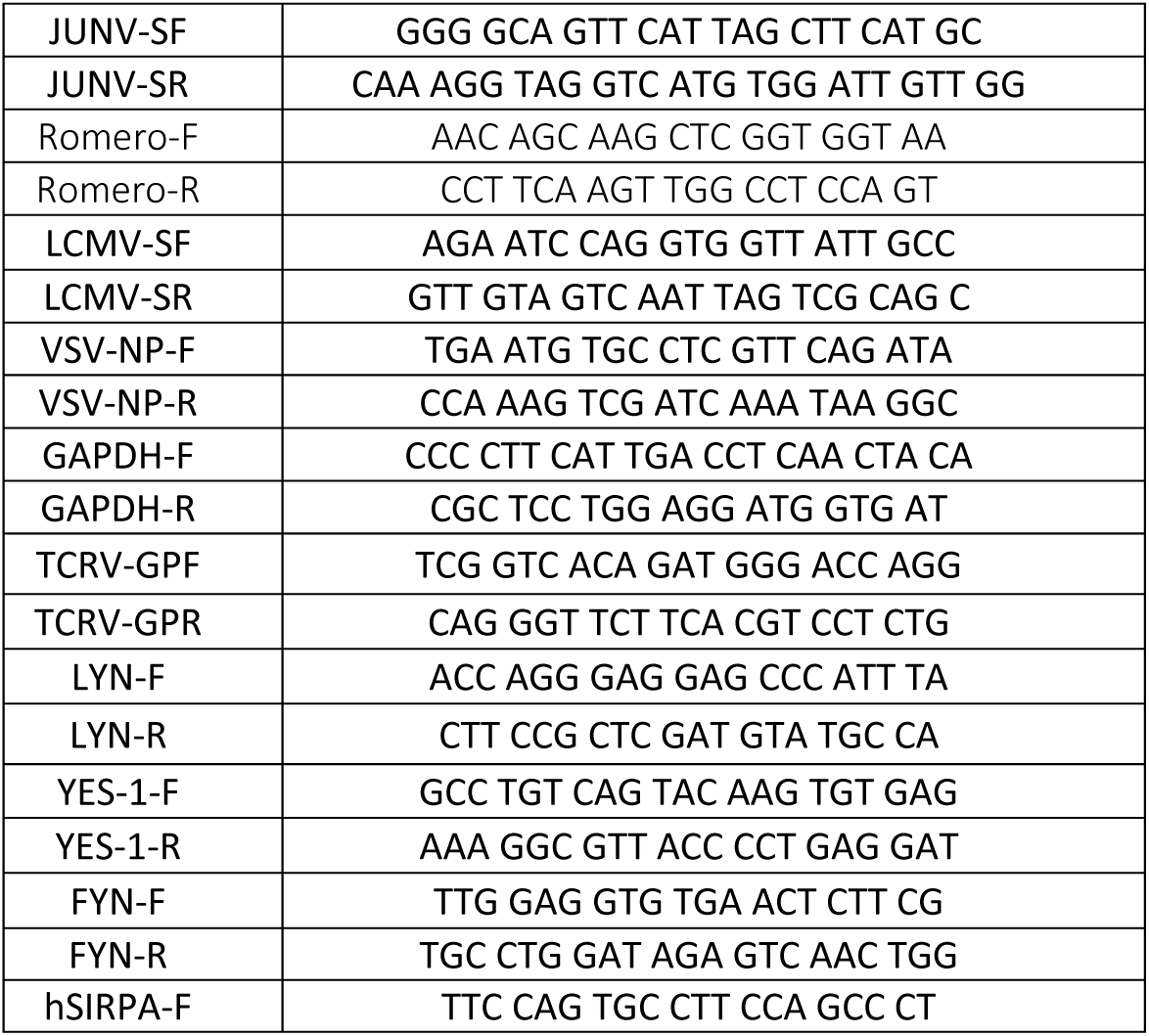
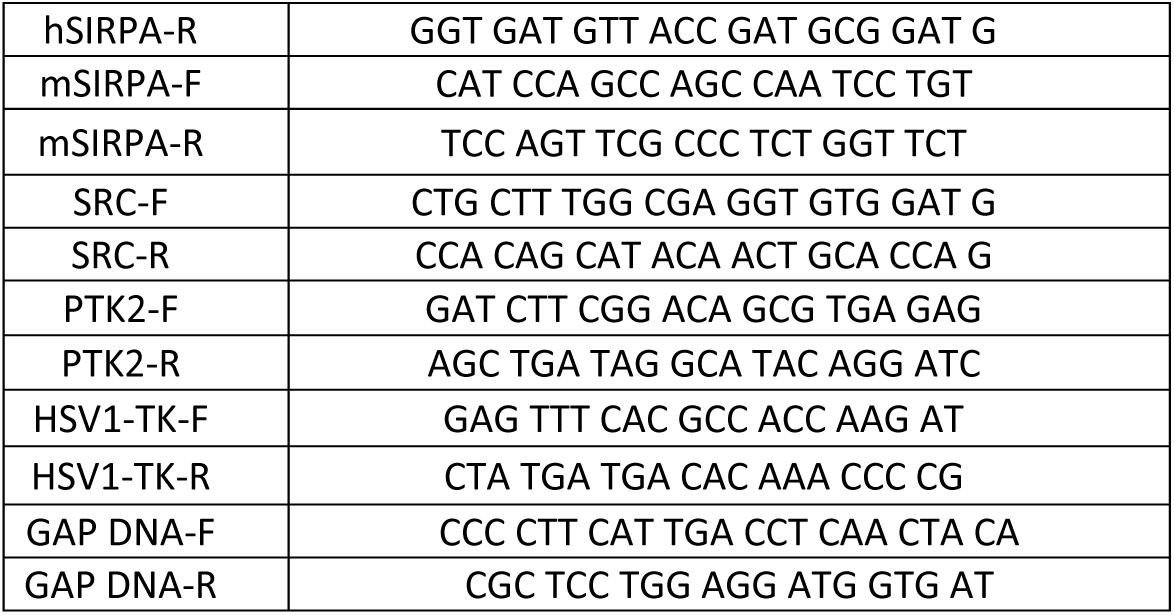
PCR primers.

### Generation of mouse primary cells

Ear fibroblasts from 8– to 12-week-old SIRPA KO and WT mice were cultured in DMEM complete and infected as previously described (Sarute et al., 2021).

### shRNA knockdown cells

To create shCTRL and shSIRPA cells, Mission® shRNA lentivirus transduction particles targeting SIRPA (Sigma Aldrich) were used to transduce U2OS cells, and the cells were selected in puromycin (1 μg/ml) (Sigma). Cell fractionation was carried out with a kit as recommended by the manufacturer (Thermo Scientific). The level of knockdown was confirmed in the cytoplasmic, membrane, and total fractions by immunoblotting with anti-SIRPA antibodies (Cell Signaling Technology).

### Viruses

JUNV-C1 (obtained from the WRCEVA) and VSV-eGFP were propagated in Vero E6 cells; LCMV (Armstrong strain) and Tacaribe virus (TRVL-11573; BEI Resources) were propagated in BHK-21 cells. Viruses were propagated as previously described (Sarute et al., 2021). Briefly, cells were infected at 70% confluence at an MOI of 0.01 for 24 hours. Cells were then washed with phosphate-buffered saline (1X PBS) and refilled with DMEM with 2% FBS. Virus supernatants were collected at day 3–5 dpi (LCMV), day 5–8 dpi (JUNV-C1), or day 7–10 dpi (TCRV, VSV) and purified through ultracentrifugation with a 30% sucrose cushion. Purified viruses were then resuspended in 1X PBS, aliquoted, and stored at –80°C. HSV-1 (KOS strain; ATCC VR1493) was grown on Vero E6 cells and propagated as previously described (REF). ZIKV strain PF13 was obtained from Michael Diamond, Washington University. Viral stocks were propagated in C6/36 cells and culture supernatants were harvested at 6 dpi. The molecular clone of the Romero strain of JUNV was rescued in BHK-21 cells as previously described and kindly provided by Slobodan Paessler (University of Texas Medical Branch) (Emonet et al., 2010). The stock used was prepared from a single passage in Vero cells. All JUNV-Romero experiments were performed at Utah State University under Select Agent-approved, BSL-3+ containment.

### Virus titration

JUNV titers were determined by infectious center assays, TCRV was titrated by TCID50, and LCMV, VSV, and HSV-1 titers were determined by plaque assays in Vero E6 cells, as previously described (Emonet et al., 2010; Sarute et al., 2021).

### Plasmids

The H5N1, MACV GP, VSV G, and SIRPA expression plasmids were previously described (Sarute et al., 2021; Sarute et al., 2019). The SHP2, SRC, FYN and LYN expression plasmids were obtained from Addgene (8382, 42202, 74509 and 35958, respectively) (Chougule et al., 2016; Luttrell et al., 1999; Yoo et al., 2011). When different amounts of plasmids were transfected, empty vector (EV) plasmid [pcDNA 3.1/Myc (+)] was added to keep the total amount of transfected DNA equal for all samples.

### Pseudoviruses

H5N1-HIV, MACV-HIV, and VSV-HIV pseudoviruses were produced by co-transfection of the replication-defective HIV vector (pNL4-3.Luc.RE) and plasmids for the H5N1 A, MACV GP, or VSV G proteins into 293T cells, using a polyethylenimine-based transfection protocol, as described previously (Hussein et al., 2020; Tisoncik et al., 2011). Briefly, 6 hours after transfection, cells were washed with 1X PBS, refilled with fresh media for 24 hours. After 24 hours, virus supernatants were collected and purified by filtering through a 0.45 μm filter (Sigma).

### *In vitro* and ex *vivo* infections

U2OS cells were infected with viruses as indicated in the figure legends, and viral RNA or DNA levels were determined by RT-qPCR or qPCR, respectively. U2OS or A549 cells were infected with pseudoviruses for 48 hours, and luciferase activity was quantified using the Luciferase Assay System (Promega). THP-1 cells were infected with infectious viruses at an MOI of 1, and the viral RNA levels were determined by RT-qPCR. Primary fibroblasts were infected as previously described (Sarute et al., 2021).

### Antibodies

FLAG and Myc-tagged SIRPA constructs were detected using either a mouse anti-FLAG (M2) (Sigma #F1804) or a mouse anti-Myc (CST #2276S) antibody. Endogenous SIRPA was detected using a rabbit polyclonal antibody (CST #13379S). HA-tagged SHP2 construct was detected using a rabbit anti-HA (CST #3724S). Endogenous SHP2 and phospho-SHP2 (Y542) were detected using a rabbit monoclonal antibody (CST #3397S) and (Abcam #AB62322), respectively. Tyrosine-phosphorylated proteins were detected with a mouse monoclonal antibody (Sigma #05-321X). Src-family kinases SRC, FYN, and LYN were detected with rabbit monoclonal antibodies (CST #2109S), mouse monoclonal antibody (Santa Cruz #sc-365913), and mouse monoclonal antibody (Santa Cruz #sc-7274), respectively. FAK and phospho-FAK (Y397) were detected with rabbit polyclonal antibodies (CST #3285S) and (CST #3283S), respectively. glyceraldehyde-3-phosphate dehydrogenase (GAPDH) and β-actin served as internal controls and were detected using a rabbit polyclonal (CST #2118L) and a mouse monoclonal (Proteintech #66009-1-Ig), respectively.

### Western blots and Immunoprecipitation

Cells were lysed with RIPA buffer containing Halt Protease and Phosphatase Inhibitor Cocktail (Thermo Fisher #78442). Protein extracts were sonicated on ice, quantified by Bradford assay (BioRad), resolved on 4–15% gradient SDS-polyacrylamide gels, and transferred to polyvinylidene difluoride (PVDF) membranes (Millipore). Membranes were blocked and probed in 3% BSA (TBST). For immunoprecipitation, protein extracts were incubated with primary antibody and Protein A/G PLUS-Agarose (Santa Cruz #sc-2003) overnight, followed by three washes, resolved on 4–15% gradient gels and transferred to PVDF membranes. FLAG and Myc-tagged SIRPA constructs were pulled down using mouse anti-FLAG (M2) (Sigma #F1804) or a mouse anti-Myc (CST #2276S) antibody, respectively. Endogenous SIRPA was pulled down by a rabbit polyclonal antibody (LSBio #LS-C353657). HA-tagged SHP2 was pulled down using a rabbit anti-HA (CST #3724S).

### Deglycosylation

SIRPA was deglycosylated using the Protein Deglycosylation Mix II (NEB #P6044S) according to the manufacturer. After deglycosylation, protein extracts were incubated with anti-SIRPA antibody and Protein A/G overnight, followed by three washes, resolved on 4–15% gradient gels and transferred to PVDF membranes.

### PLAs

U2OS cells were seeded on glass coverslips in 24-well plates, co-transfected with SIRPA-Flag and SHP2-HA or pcDNA control plasmids for 24 hours, bound to TCRV on ice for 1 hour (MOI = 5), warmed for for 5 or 15 minutes at 37°C to allow internalization, and fixed and permeabilized with ice-cold methanol for 15 minutes. Blocking and staining were performed with NavinciFlex 100 MR PLA probes and detection reagents (Navinci). Images were taken at 40X on a Keyence BZ-X710 microscope and analyzed with the BZ-X analyzer. Cells that showed red dots (indicating colocalization) were considered positives, and the number of positive cells in each picture was used to calculate percent positivity. At least eight fields per coverslip were imaged. The primary antibodies used were rabbit anti-DYKDDDDK Tag (Cell Signaling Technology) and mouse anti-HA tag (Cell Signaling Technology).

### *In vivo* infections

Four-week-old huTfR1 Tg mice were weighed 1 day before infection, and equal numbers of males and females were assigned to groups (n = 8 per group) to minimize weight and age differences. Mice were treated IP with 500 µg of MAR1-5A3 antibody (Lot # 1025L200; Leinco Technologies) 24 hours before infection. BTT-3033 (1 mg/kg) was administered IP beginning at 1 hour pre-infection and continued once daily (q.d.) for 6 days. Favipiravir (200 mg/kg/day) was administered IP twice daily, starting 1 hour before virus challenge and continuing through the morning of day 6 post-infection, when the mice were sacrificed; serum and spleen were collected for viral titer determination.

### Nucleic acid isolation and RT-qPCR

DNA was isolated with the DNeasy Blood & Tissue Kit (Qiagen). Total RNA was isolated using the RNeasy kit (Qiagen) for cell cultures and tissues. RNA was reverse transcribed using the SuperScript III First-Strand Synthesis System (Invitrogen) according to the manufacturer’s protocol. Viral and cellular RNAs were detected by RT-qPCR using a QuantStudio 5 Real-Time PCR System (Applied Biosystems) with specific primer pairs (Table 1). Target RNA was normalized to GAPDH. RT-qPCR reactions were performed using Power SYBR Green Master Mix (Applied Biosystems). The amplification conditions were as follows: 50°C for 2 min, 95°C for 10 min, and 40 cycles of 95°C for 15 s and 60°C for 1 min. The efficiency of amplification was determined for each primer set by a standard curve with 10-fold serial dilutions of DNA of known concentration. The slope values of the standard curves for the primer pair amplicons ranged from 3.5 to 3.2. For each PCR reaction, a no template control was included, and each sample was run in triplicate. For detection of JUNV-Romero, all RT-qPCR reactions were performed using QuantStudio™ 5 Real-Time PCR System and SensiFAST™ Probe No-ROX One-Step Kit with the following amplification cycle conditions: reverse transcription at 45 °C for 10 min, polymerase activation at 95 °C for 2 min followed by 40 cycles of 95 °C for 5 s and 60 °C for 20 s. Primers and probe targeting N nucleocapsid protein of JUNV were used (Table 1). The probe sequence was 5’– /56-FAM/GCA GTC TAG CAG AGC ACA GTG TGG /3BHQ_1/ –3’. A commercial primer/probe set for GAPDH (Hs.PT.39a.22214836) was used as reference control for data normalization. Data were analyzed and expressed as fold change using the method described by (Pfaffl, 2001).

### VIAs

Cells were incubated with JUNV-C1, TCRV, LCMV or VSV (MOI = 5) on ice for 1 hr, shifted to 37°C for 60 min, treated with 1 mg/ml of Proteinase K (Gibco) for 45 min on ice to remove virus bound to the cell, followed by treatment with 2 mM of phenylmethylsulfonyl fluoride (Sigma) to inactivate the Proteinase K. RNA was isolated and used for RT-qPCR.

### Chemicals and biologicals

BTT-3033 and FAK inhibitor 14 (FIB-14) were purchased from TOCRIS, LHF-535 and favipiravir (6-fluoro-3-hydroxy-2-pyrazinecarboxamide) from TargetMol Chemicals and PP1 from Cayman Chemical. Inhibitors were reconstituted in DMSO according to the manufacturer’s instructions. Cells were pre-treated with inhibitors or DMSO for 1-2 hours (time and concentrations indicated in the figure legend), incubated with virus with inhibitors for 1 hour at 37°C, washed and cultured in fresh media for 24 hours, and harvested for RNA. For VIAs, cells were pre-treated with inhibitors or DMSO for 1-2 hours, incubated with virus with inhibitors for 1 hour on ice then at 37°C in the presence of inhibitor for 1 hour, followed by Proteinase K and PMSF treatment, and harvested for RNA. For pseudovirus infection, cells were treated with inhibitors or DMSO for 1 hour, inoculated with pseudovirus with inhibitors for 1 hour at 37°C, cultured in fresh media for 48 hours, and harvested for luciferase activity. Cells treated with FAK inhibitor 14 and PP1 at different concentrations were run analyzed by western blots to check the inhibition of FAK (Y397) and SIRPA phosphorylation.

### Plasmid transfection

Plasmids were transfected into 80–90% confluent U2OS cells seeded in 6-well or 24-well plates for 24 hr using Lipofectamine 2000 reagent (Thermo Scientific), according to the manufacturers’ instructions. After 24 hours, cells were washed with 1X PBS and downstream experiments were carried out. Transfection efficiencies were confirmed by western blots.

### FACS analysis

Adherent cells were dispersed with 1X PBS/1mM EDTA and stained with FITC anti-human SIRPA (CD172a; eBioscience). The cells were stained and washed with FACS buffer (1% FBS 0.01% sodium azide in 1X PBS) and were analyzed in a BD LSRFortessa cell analyzer (BD Biosciences) using FlowJo v10 software (Tree Star, Inc.).

### Bead phagocytosis assay

THP1 cells seeded in 24 well plates, differentiated with 30ng/ml PMA for 24 hours and then activated with IL-4 (100 ng/ml) for 48 hours. Cells were pretreated with 20μM BTT-3033 (Tocris) in media for 1 hour at 37°C. FluoSphere carboxylate beads (1 μm/yellow-green; Invitrogen #F8823) were added with 20μM BTT-3033 in media for 1 hour at 37°C (200 beads per cell). Cells were washed thrice with ice cold 1X PBS. Cells were flushed in 1X PBS to dislodge them and FACS was run using the BD LSR Fortessa with HTS (FITC filter) and analyzed using FlowJo software. Fifty μl of 0.4% Trypan blue (Molecular Probes #T10282) per 500 μl of sample was added just before running FACS to quench beads that were not phagocytosed.

### Statistical analysis

Each experiment was done with 3 technical replicates per experiment. Data shown is the average of at least 3 independent experiments, or as indicated in the figure legends. Statistical analysis was performed using GraphPad 8.1/PRISM software. Tests used to determine significance are indicated in the figure legends.

### Data availability statement

All raw data have been deposited in the Mendeley data set found at 10.17632/2vz4gvvvyv.1.

## Acknowledgements

We thank David Ryan for help with mice and Justin Richner for help with the ZIKV infections. Viruses and cells were obtained from the WRCEVA (JUNV C1), Michaela Gack and Sean Whelan (VSV-eGFP), Michael Diamond (ZIKV strain PF13) and from BEI Resources (TCRV and NR9456 cells), and the H5N1 expression plasmid was obtained from Lijun Rong. Supported by NIH/NIAID 5R01AI159290. C. T.-H. was supported by NIH/HL 2T32HL007829.

## Supplementary Fig. Legends

**Figure S1.** SIRPA and SHP2 interaction depends on tyrosine phosphorylation in ITIM region and virus induces SIRPA phosphorylation. A) U2OS cells were co-transfected with WT SIRPA (Flag-tagged), Δcyto or 1-4A expression vectors along with a SHP2 (HA-tagged) expression vector. PLA using anti-Flag antibody and anti-HA antibody was carried out. Shown to the right of the photographs is quantification of the appearance of the PLA+ spots (average ± SD of 3 independent experiments). For each experimental condition, at least eight fields were imaged. Unpaired T tests were used to determine significance. ns, not significant, ***, P ≤ 0.0003; *, P ≤ 0.01. B) SIRPA deglycosylation confirmation. shControl or shSIRPA cells were lysed for de-glycosylation or mock treatment. De-glycosylated cell lysates were immunoprecipitated with anti-SIRPA antibody, and western blots were probed with anti-phospho-tyrosine or –SIRPA antibodies. The input blot was stripped and probed with anti-GAPDH as a loading control. Red arrow, glycosylated SIRPA; blue arrow, deglycosylated SIRPA. C) Incubation of JUNV C1 with mouse macrophages leads to increased SIRPA phosphorylation that is inhibited by PP1. NR9456 cells were incubated with JUNV C1 for 15 min. in the presence or absence of PP1. Cell lysates were immunoprecipitated with anti-SIRPA antibody, and western blots were probed with anti-phospho-tyrosine antibody. Abbreviations: Veh, vehicle.

**Figure S2.** Src-family kinases control tyrosine phosphorylation of SIRPA. A) U2OS cells were co-transfected with SIRPA and the indicated SFK expression plasmids. The cell lysates were immunoprecipitated with anti-SIRPA antibody, and western blots were subjected to probing with the indicated antibodies. B) U2OS cells were co-transfected with the indicated SIRPA (Flag-and Myc-tagged) and FYN (Flag-tagged) plasmids. The cell lysates were immunoprecipitated with anti-Myc antibody, and western blots were probed with anti-FYN or –SIRPA antibodies. C) U2OS cells were transfected with indicated Src-family kinase plasmids. Cells were lysed for de-glycosylation and the deglycosylated cell lysates were immunoprecipitated with anti-SIRPA antibody. Western blots were probed with the indicated antibodies. D) U2OS cells were transfected with indicated siRNA. Cells were lysed for deglycosylation and the de-glycosylated cell lysates were immunoprecipitated with anti-SIRPA antibody, and western blots were probed with the indicated antibodies. Quantification of phospho-tyrosine level normalized to SIRPA level using Image Lab are shown in the bar graph. Shown is the average ± SD of 3 independent experiments. One-way ANOVA was used to determine significance. *, *P* ≤ 0.05; ***, *P* ≤ 0.001.

**Figure S3.** Integrin inhibitor BTT-3033 inhibits JUNV Romero and ZIKV infection. A) BTT-3033 has limited toxicity with U20S cells up to 10 μM concentrations with 3 hr incubation. B) U2OS shSIRPA cells were treated with BTT-3033 (10μM) or LHF-535 (20nM) and infected with JUNV Romero. RNA was isolated at 24 hpi and viral RNA was measured by qPCR. Shown are 2 independent experiments for each drug, with triplicate technical replicates in each experiment. C) U2OS cells were treated with BTT-3033 and infected with ZIKV. RNA was isolated at 24 hpi and viral RNA was measured by qPCR. Shown is the average ± SD of 3 independent experiments. An unpaired T test was used to determine significance. ****, P ≤ 0.001.

**Figure S4.** Knockdown of FAK or ITGA5 decreases SIRPA expression and increases virus infection. A) U2OS transfected with FAK siRNAs were infected with the indicated viruses. RNA was isolated at 14 (VSV) or 24 (JUNV-C1, LCMV,) hpi. Viral and FAK nucleic acids were measured by RT-qPCR. Shown is the average ± SD of 3 independent experiments. Student’s T test was used to determine significance. *, *P* ≤ 0.02; **, *P* ≤ 0.001; ****, *P* ≤ 0.0001. B) U2OS cells were transfected with FAK or ITGA5 siRNAS and at 48 hr post-transfection, SIRPA RNA levels were measured. Shown is the average ± SD of 3 independent experiments. One-way ANOVA was used to determine significance. ****, *P* ≤ 0.0001. C) U2OS cells were treated with indicated concentrations of FIB-14. SIRPA surface expression was analyzed by FACS using antibody to the ectodomain.

**Figure S5.** Phagocytosis is inhibited by BTT-3033 treatment but is not affected by SIRPA levels. A) SIRPA surface levels in WT (dark orange) and shSIRPA-THP1 (light orange) cells. B) BTT-3033 diminishes bead phagocytosis by THP1 cells. C) SIRPA knockdown in THP1 cells does not affect bead internalization. Shown to the right of the histogram are the mean and standard deviation of two experiments. Pink: shSIRPA-THP1; grey: WT THP1 cells.

