## Supplementary figures and images for "SIRPA suppresses integrin-dependent virus endocytosis"

### Supplemental FIgures

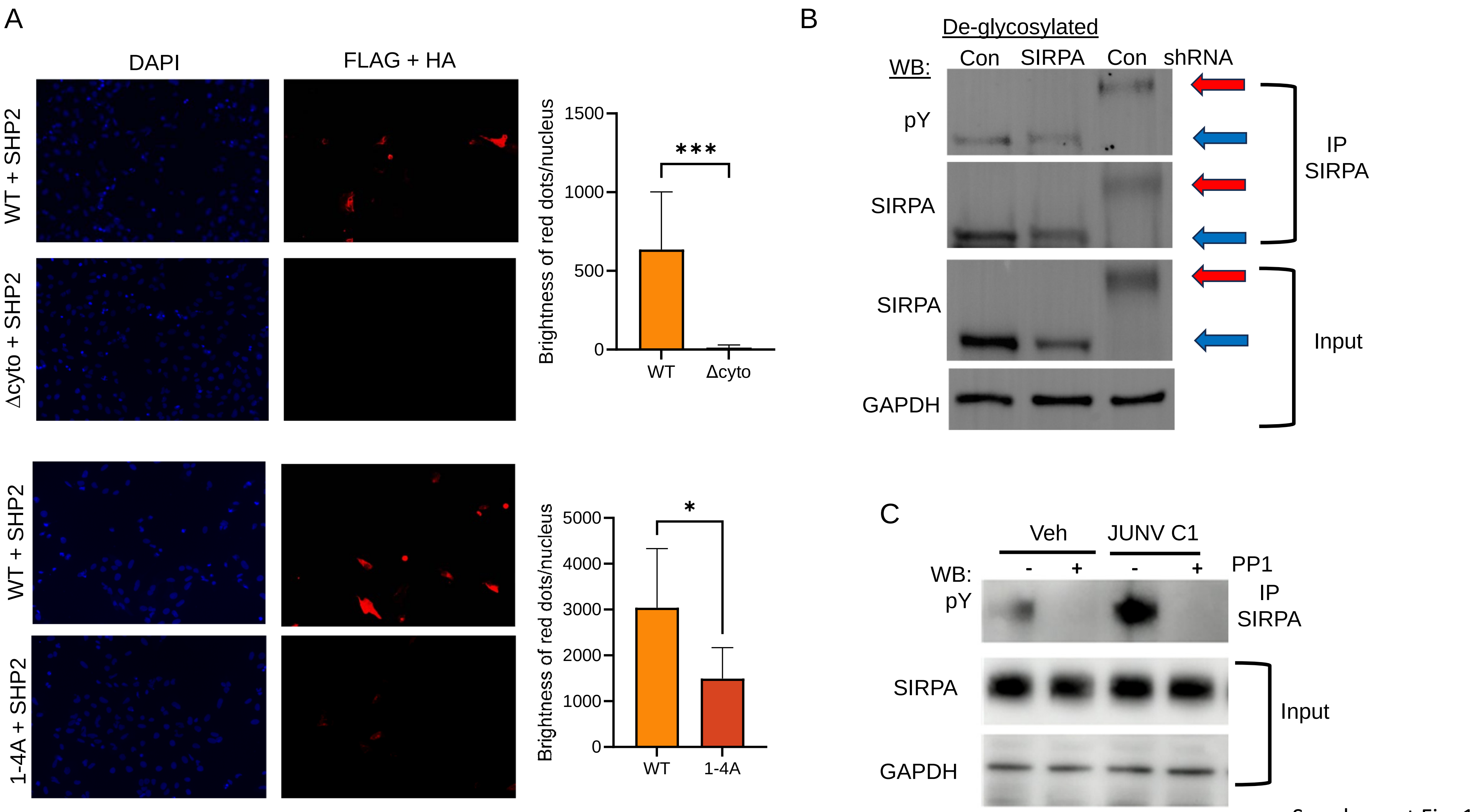


Supplement Fig. 1

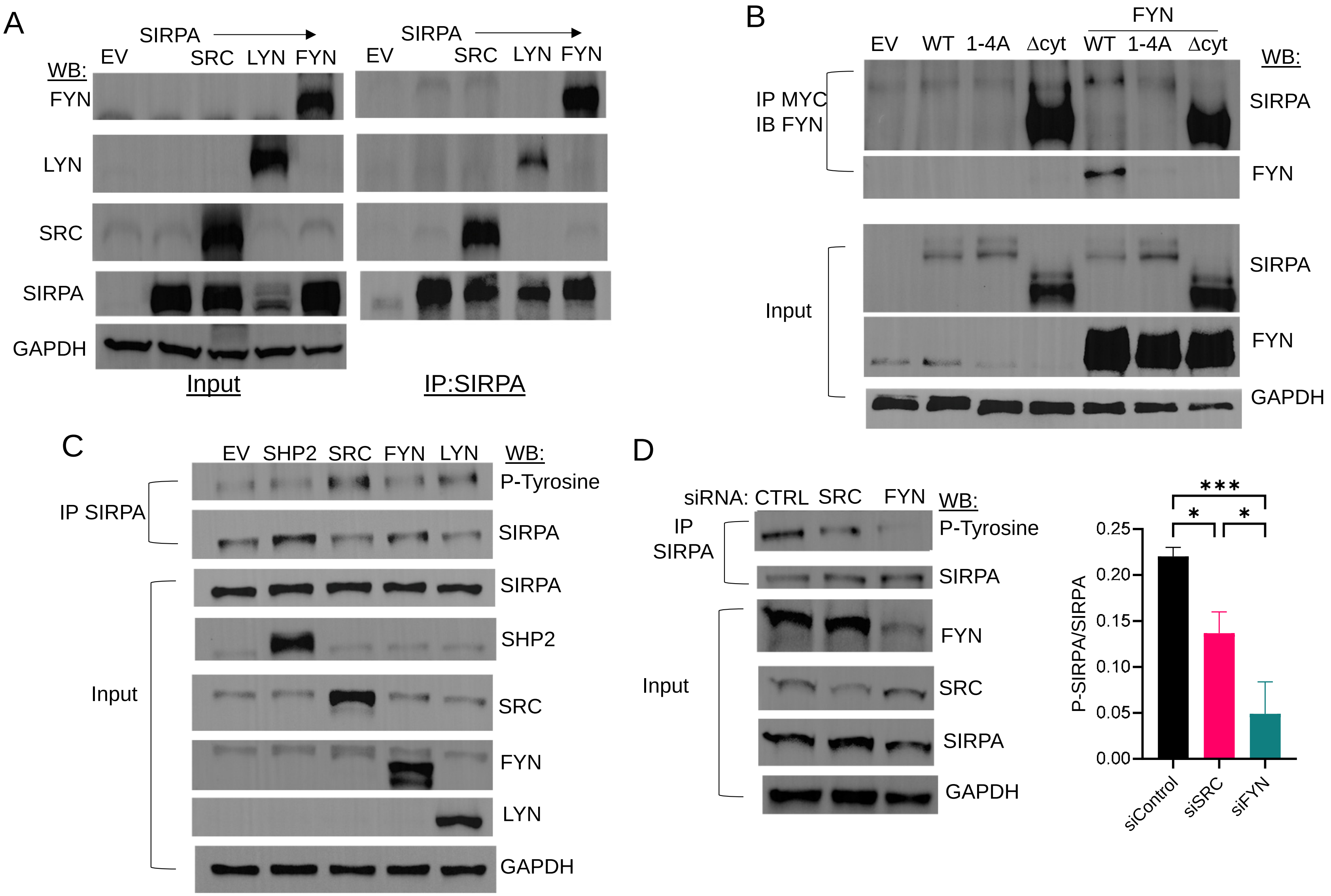


Supplement Fig. 2

A

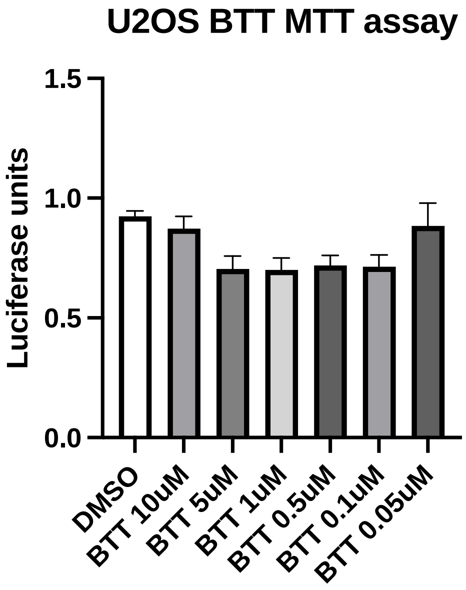


B

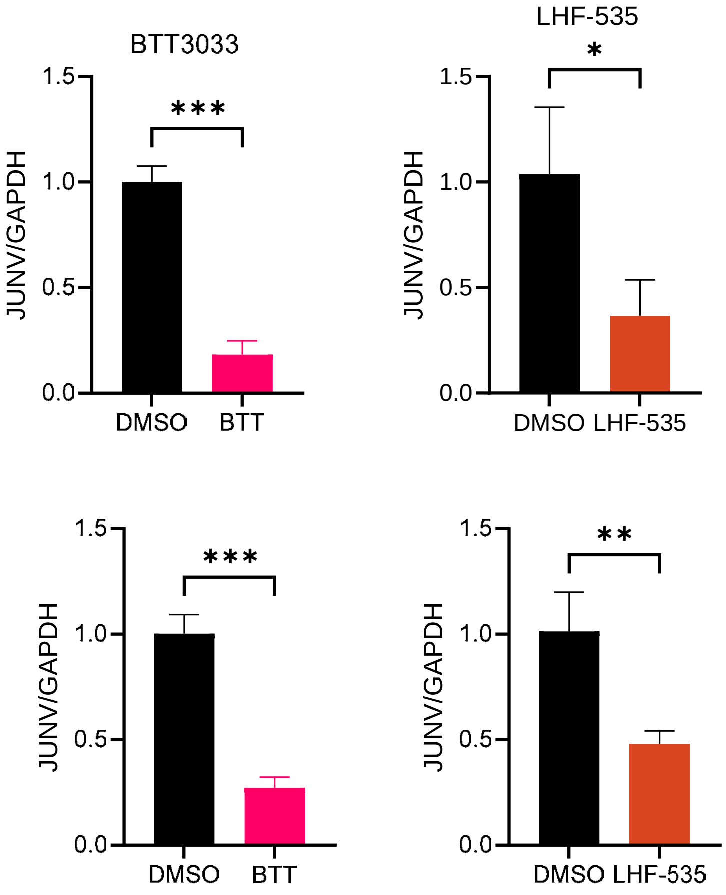


C

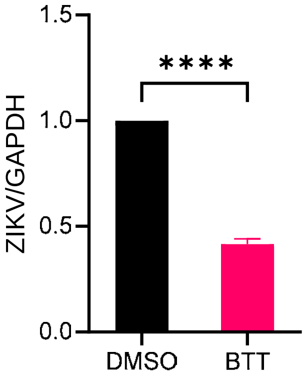


Supplementary Fig. 3

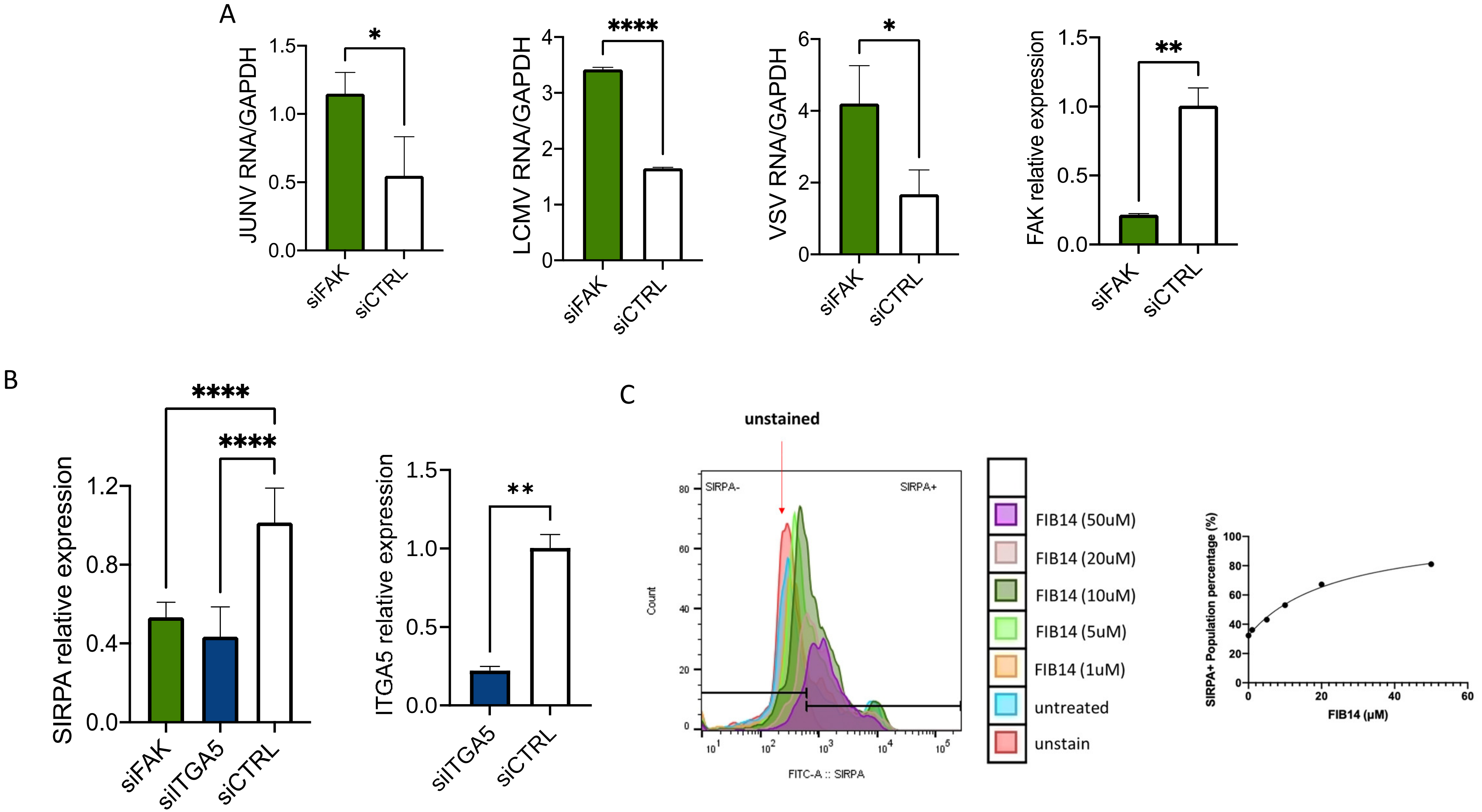


Supplementary Fig. 4

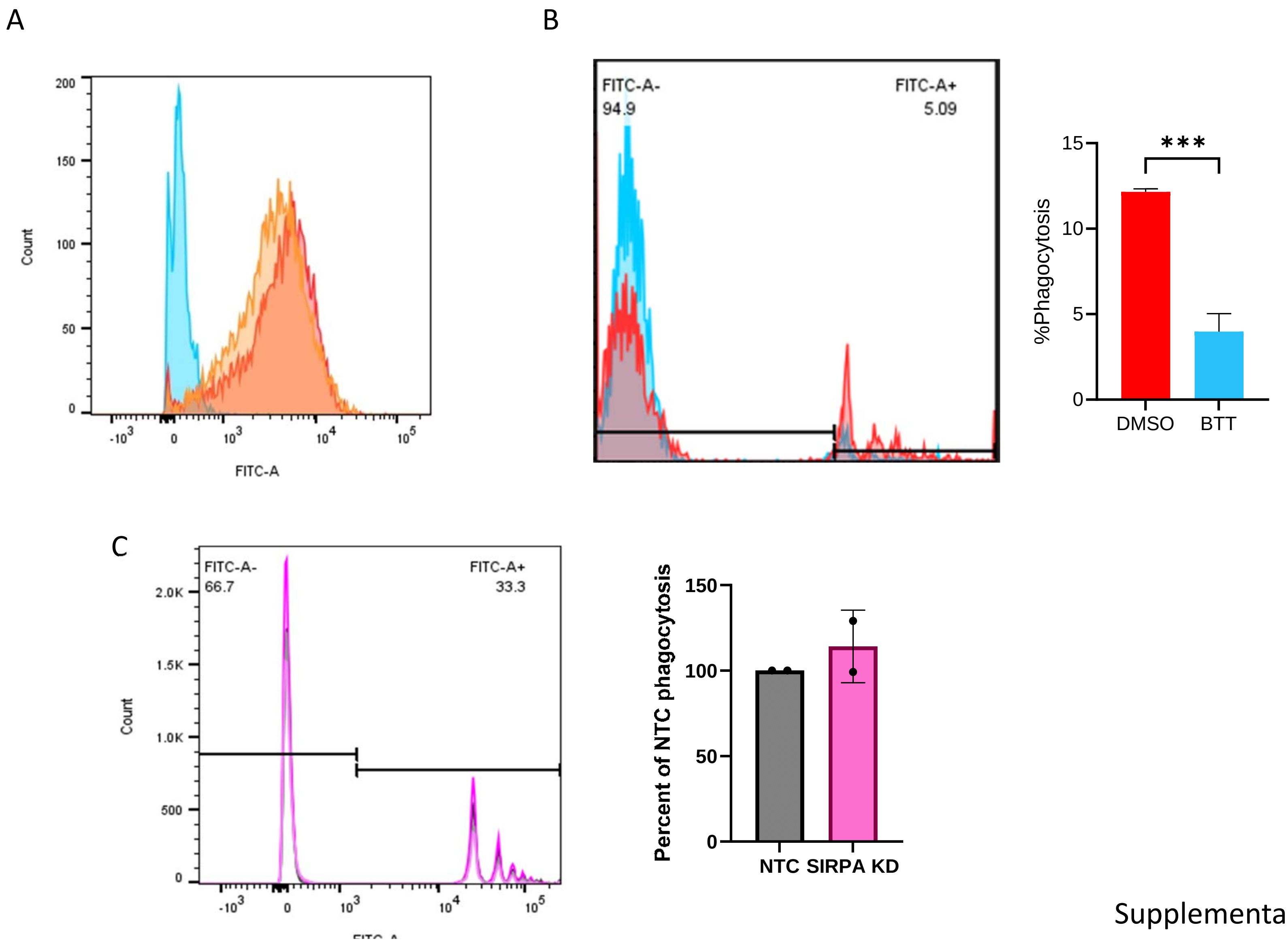


Supplementary Fig. 5
